# Fungal mycelia and bacterial thiamine establish a mutualistic growth mechanism

**DOI:** 10.1101/2020.07.12.199612

**Authors:** Gayan Abeysinghe, Momoka Kuchira, Gamon Kudo, Shunsuke Masuo, Akihiro Ninomiya, Kohei Takahashi, Andrew S. Utada, Daisuke Hagiwara, Nobuhiko Nomura, Naoki Takaya, Nozomu Obana, Norio Takeshita

**Affiliations:** Microbiology Research Center for Sustainability (MiCS), Faculty of Life and Environmental Sciences, University of Tsukuba, 305-8572, Japan

**Author notes:** Correspondence to: Nozomu Obana, Norio Takeshita. Equal contribution (Gayan Abeysinghe, Momoka Kuchira). MiCS and Transborder Medical Research Center, Faculty of Medicine, University of Tsukuba (Nozomu Obana).

**Keywords:** fungal highway, bacterial dispersal, thiamine, mutualism

## Abstract

Physical spaces and nutrients are prerequisites to the survival of organisms while few interspecies mutual strategies are documented that satisfies them. Here we discovered a mutualistic mechanism between filamentous fungus and bacterium, *Aspergillus nidulans* and *Bacillus subtilis*. The bacterial cells co-cultured with the fungus traveled along mycelia depending on their flagella and dispersed farther with the expansion of fungal colony, indicating that the fungal mycelia supply space for bacteria to migrate, disperse and proliferate. Transcriptomic, genetic, molecular mass and imaging analyses demonstrated that the bacteria reach the mycelial edge and supply thiamine to the growing hyphae, resulting in a promotion of hyphal growth. The thiamine transfer from bacteria to the thiamine non-auxotrophic fungus is directly demonstrated by stable isotope labeling. The simultaneous spatial and metabolic interactions demonstrated in this study, reveal a mutualism that facilitates the communicating fungal and bacterial species to obtain environmental niche and nutrient respectively.

## Introduction

Microbes ubiquitously live in nearly every ecological niche. Different species coexist in certain habitats and interact with each other. Microbes often constitute communities and share available metabolites (1). Natural auxotrophic strains grow in the presence of external nutrients which are provided by members of the local microbiota (2). Since such nutrients limit microbial growth, acquiring them within communities is essential for auxotrophs to utilize an ecological niche.

Bacteria and fungi comprise a large fraction of the biomass in soil (3, 4). Since they interact with each other to carry out their characteristic functions in the ecosystem, a better knowledge of bacterial-fungal interactions is important for understanding the microbial ecosystem, which is closely related to agriculture, medicine, and the environment (5). Interkingdom interactions are driven by diverse factors such as antibiotics, signaling molecules, cooperative metabolism, and physical interactions (6). In certain scenarios, bacteria physically attach to fungal tube-shaped hyphal cells thus enabling changes in their metabolism either antagonistically or beneficially (7). It has been shown that fungal hyphae transfer nutrients and water to activate bacteria (8), while bacteria are able to induce the expression of transcriptionally inactive genes for synthesizing fungal secondary metabolites (9).

Filamentous fungi grow by extension of hyphae at their tips, thereby forming multicellular networks with branching cells at subapical regions (10). Mycelial network spreads on solid surfaces, that allows the fungus to reach spatial niches in the ecosystem. In contrast, bacteria are unicellular organisms, some of which are motile, enabling them to explore the environment in search of better spatial and nutrient conditions (11). While motility is efficient in liquid (12), bacteria can disperse farther in water-unsaturated conditions by traveling along fungal hyphal “highways” (13, 14). This interaction is considered as commensal because the fungi do not benefit from providing a “highway” for bacteria.

Here, we describe a mutualistic growth mechanism between models of filamentous fungus and gram-positive bacterium, *Aspergillus nidulans* (9, 15, 16) and *Bacillus subtilis* (17), where both organisms benefit from the fungal highways and the sharing of one vitamin, respectively.

## Results

### Bacterial movement along fungal hyphae

We tested several combinations of fungal-bacterial co-culture among our lab strains, then selected the combination of *A. nidulans* and *B. subtilis,* that are relatively common in soil, for further analysis. *B. subtilis* grew in co-culture with *A. nidulans* at the comparable rate as in liquid monoculture (Fig. S1A). Live imaging analysis showed that *B. subtilis* cells moved along the co-cultured *A. nidulans* hyphae on the agar medium (Fig. 1A, Movie S1). Some bacteria remained attached to the hyphae, while others moved along the hyphae, often reversing course abruptly and beginning to move in the opposite direction. A heat-map of the instantaneous velocity was constructed by tracking the positions of each moving cell; the results indicated weak oscillations in the instantaneous velocity over time (Fig. 1B, C and Fig. S2). Kymographs indicated that the bacterial cells moved at an average velocity of ~30 μm s^-1^ in both directions (Fig. 1D, E and Movie S2). Rapid movements were not observed in bacterial monoculture on solid medium (Fig. S3A) and were comparable to *B. subtilis* movement in fungus-free liquid medium (18). The numbers and density of motile and non-motile bacteria were not uniform on the mycelium. Some bacterial cells reached the hyphal tips, and then reversed their direction after remaining at the tip for some time (Fig. 1F and Movie S3). Other hyphae were surrounded by moving bacterial aggregates (Fig. S3B, Movie S4). The co-cultures were observed by a scanning electron microscope (Fig. 1G).

**Fig. 1.**
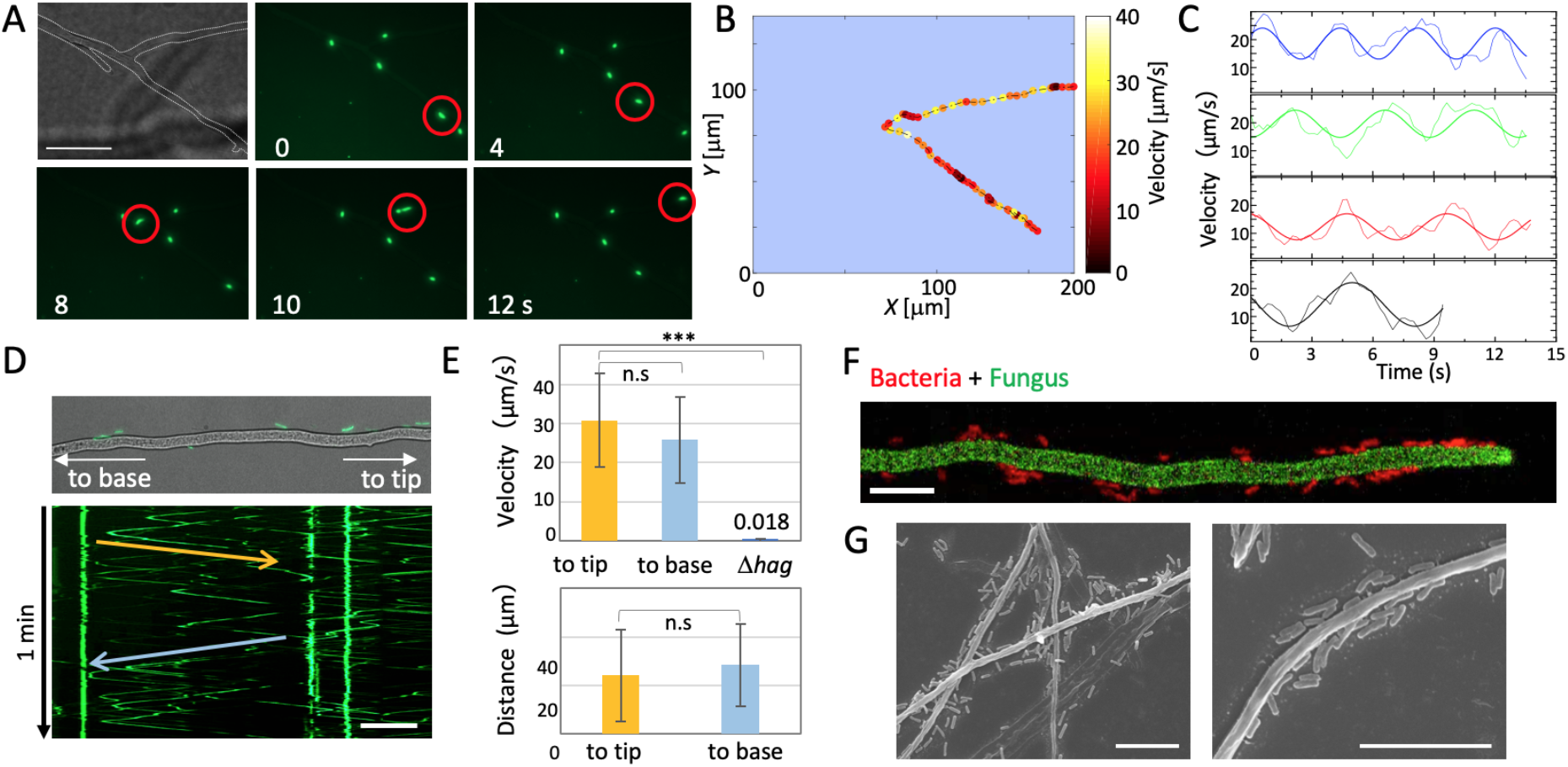
Bacterial movement along fungal hyphae. (A) Time-lapse images of *B. subtilis* (green) movement along *A. nidulans* hyphae (dotted line) for 30 s on agar media from Movie S1. Scale bar: 50 μm. (B) Heat-map of *B. subtilis* instantaneous velocity analyzed by tracking the position of the cell moving along the hypha in (A). (C) Weak oscillations in the instantaneous velocity over time are shown in different colors from each bacterial cell in B and Fig. S2. (D) Kymograph of *B. subtilis* movement along the hyphae (top) to the tip (yellow arrow) and base (blue arrow) Movie S2. Total 1 min. Scale bar: 50 μm. (E) Velocity and distance of *B. subtilis* (wild-type or Δ*hag*) movement along hyphae to the tip (yellow) or base (blue). Error bar: S.D., n = 26 (to tip), 28 (to base), 5 (Δ*hag*), *** P < 0.001. (F) *B. subtilis* cells (red) reach the tip of *A. nidulans* hyphae (green) from Movie S3. Scale bar: 10 μm. (G) SEM images of co-culture of *B. subtilis* and *A. nidulans.* Scale bar: 10 μm.

*B. subtilis* strain 168, which is defective in producing the biosurfactant surfactin necessary for swarming on solid agar plates (19), still moved along the hyphae. In contrast, the flagellar-deficient mutant (Δ*hag*) traveled towards the hyphal tips at considerably lower rates (0.018 μm s^-1^, 0.05% of control strain) (Fig. 1E, Fig. S3C and Movie S5), indicating that *B. subtilis* move along the hyphae using flagella.

### Bacterial dispersal on growing fungal hyphae

*B. subtilis* generated smaller colonies on the agar medium than *A. nidulans* (Fig. 2A). The size of the co-cultured *A. nidulans* colony was ~30 % larger than that of fungal monoculture. Fluorescence-tagged *B. subtilis* were observed on the colony periphery of the co-culture. This dispersal depended on bacterial movement towards the hyphal tips (Fig. 2B and Movie S6). The rate of bacterial colony expansion was 7-fold faster (154 ± 33 μm h^-1^) than that of *B. subtilis* monoculture (21 ± 3 μm h^-1^) (Fig. 2B, C and Movie S7). The flagella were indispensable for bacterial dispersal along the growing hyphae and for the cells to reach the hyphal tips (Fig. 2A-C and Movie S8). The colony expansion rate of the Δ*hag* strain was comparable to that in the wild-type monoculture (Fig. S4). Since the bacterial movement along the hyphae was much faster (approximate 30 μm s^-1^) than the extension rate of growing hyphae, bacterial cells reached the ends of the hyphal tips (Fig. 2D and Movie S9). The mycelium network appears to supply a space for bacteria to migrate, disperse and proliferate (Fig. 2E and Movie S10). Indeed, the bacterial proliferation measured by the amount of bacterial DNA, was higher in the co-culture more than the monoculture on the agar plates (Fig. S1C).

**Fig. 2.**
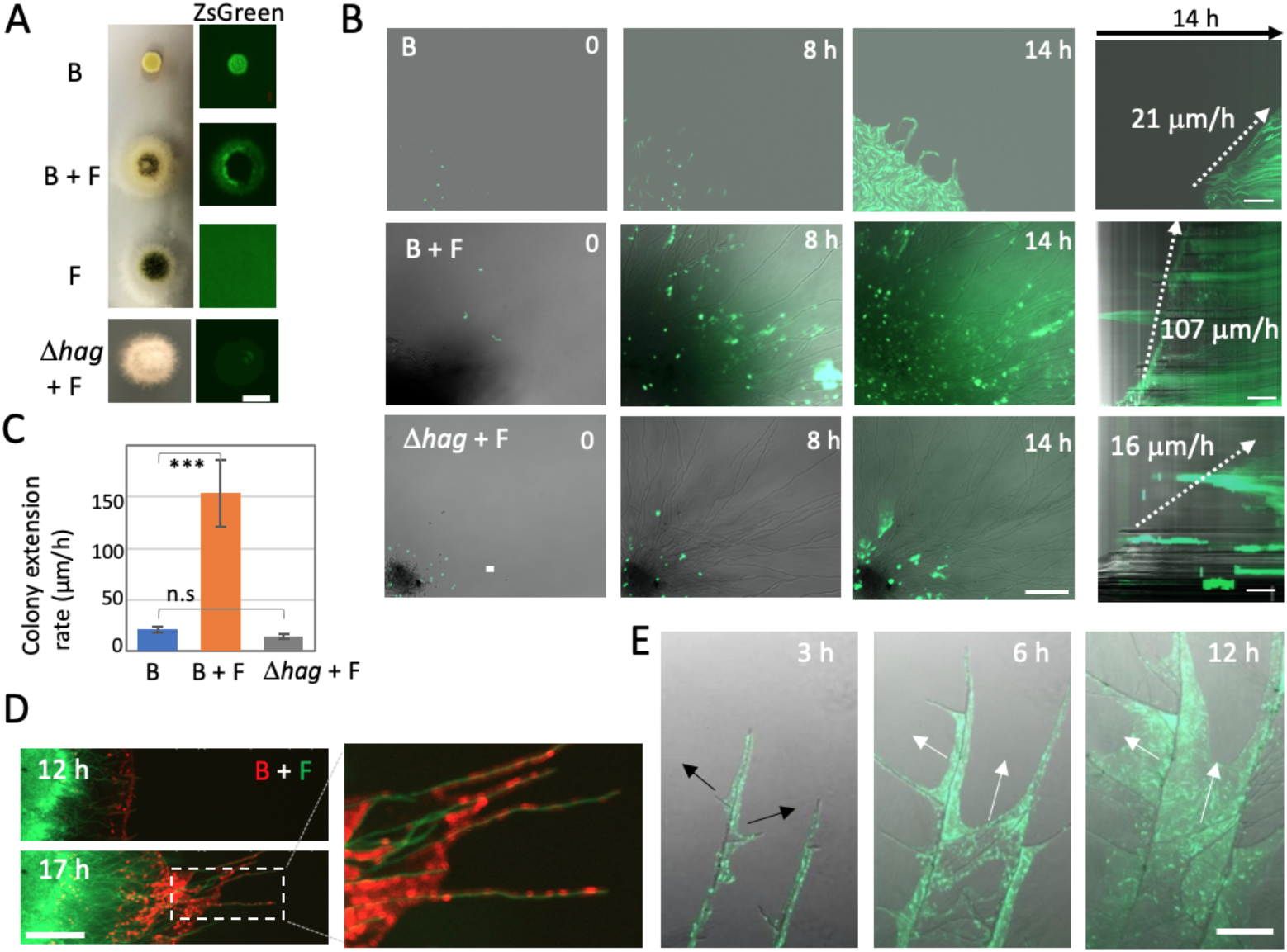
Bacterial dispersal on growing fungal hyphae. (A) Colonies of *B. subtilis* and *A. nidulans* mono-culture and co-culture, co-culture of *A. nidulans* and *B. subtilis* (Δ*hag*) (bottom). ZsGreen-labeled *B. subtilis* (right). Aerial growing hypha at the middle of colony disturb to detect the fluorescent signals in the co-culture. B: *B. subtilis,* F: *A. nidulans.* Scale bar: 500 μm. (B) Time-lapse images at 0, 8 and 14 h of *B. subtilis* mono-culture, *B. subtilis* (WT or Δ*hag*) with *A. nidulans* from Movie S6-8. Scale bar: 100 μm. Kymographs of *B. subtilis* (WT or Δ*hag*) dispersion with/without growing hyphae. Scale bar: 50 μm. (C) Colony expansion rates calculated from kymographs. Error bar: S.D., n = 5, *** P < 0.001. (D) Time-lapse *B. subtilis* (red) dispersion on growing *A. nidulans* hyphae (green) from Movie S9. Scale bar: 200 μm. (E) Time-lapse *B. subtilis* dispersion (green, white arrows) on *A. nidulans* branching hyphae (DIC, black arrows) after 17 h co-culture from Movie S10. Scale bar: 200 μm.

### Metabolic interaction through thiamine

We analyzed the effect of co-culture of *B. subtilis* and *A. nidulans* on extracellular hydrophobic metabolites (Fig. S5) and transcriptomes. RNA-sequencing analysis indicated that expression of most *B. subtilis* and *A. nidulans* genes was not affected by the co-culture. The expression of 18 genes in *B. subtilis,* including the thiamine biosynthesis operon, was induced by 2-fold in the co-culture with *A. nidulans* (Fig. 3A, Table S1 and Fig. S1D). In contrast, thiamine biosynthesis-related genes in *A. nidulans* were down-regulated in co-culture (Fig. 3A and Table S2). The upregulated genes in *A. nidulans* include asexual spore formation, nitrate inducible genes, non-ribosomal peptide synthases, and polyketide synthases (Table S3). The induction in *B. subtilis* and repression in *A. nidulans* of thiamine biosynthesis-related genes implied that the bacterium and the fungus metabolically interact *via* thiamine. We co-cultured the *A. nidulans* strain defective in thiamine biosynthesis (Δ*thiA*; putative thiazole synthase) (20) with *B. subtilis*. The Δ*thiA* fungal colony showed a severe growth defect on the plate without thiamine (Fig. 3B), which was recovered by adding thiamine. The growth defect of Δ*thiA* was recovered by co-culture with wild-type *B. subtilis*, but not by co-culture with the *B. subtilis* thiamine synthesis mutant (Δ*thi*; operon deletion) (Fig. 3B), indicating that *B. subtilis* synthesizes and supplies thiamine to *A. nidulans*.

**Fig. 3.**
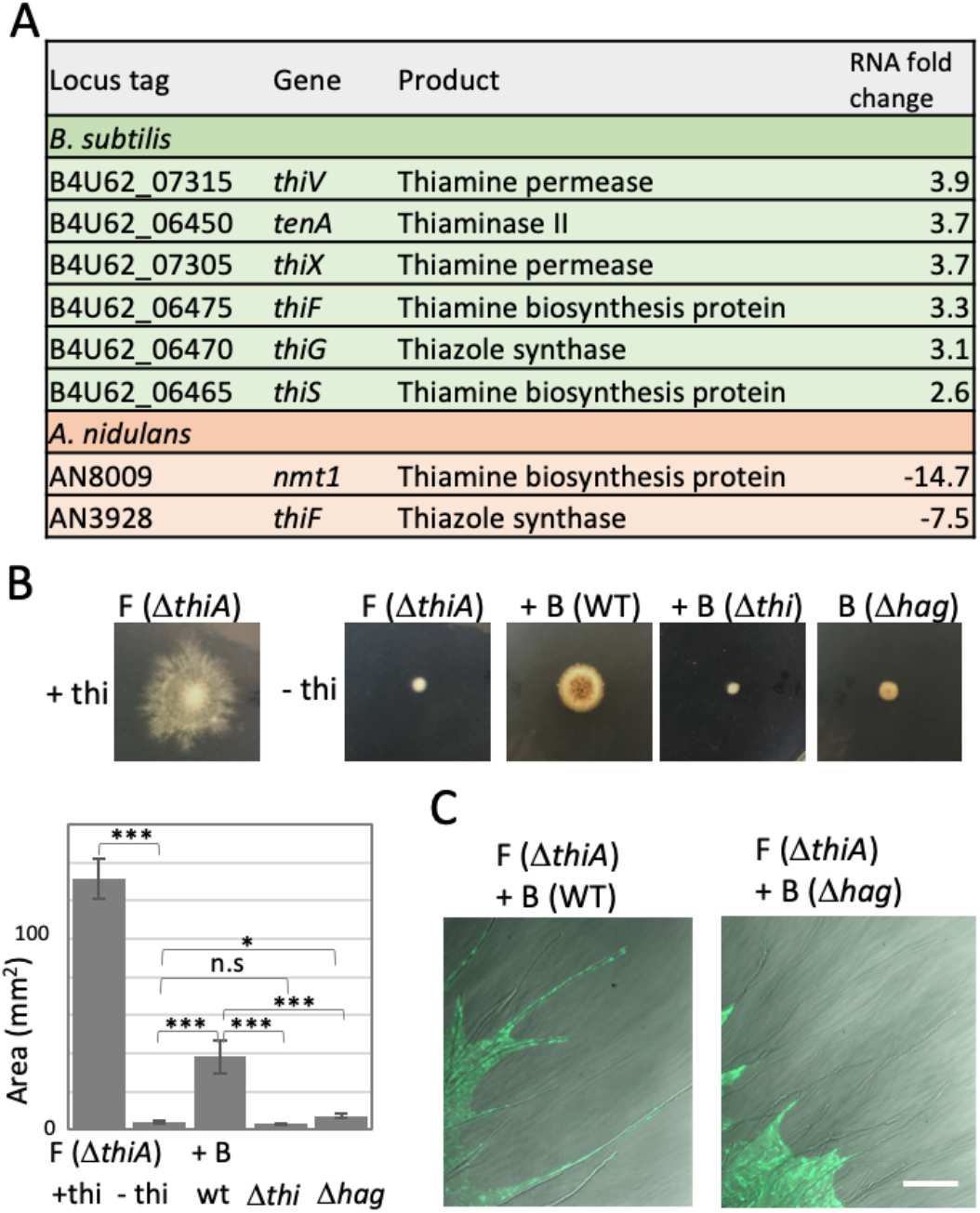
Metabolic interaction through thiamine. (A) Summary of RNA-seq analysis related to thiamine synthesis in *B. subtilis* (green) and *A. nidulans* (orange). (B) Fungal colonies of *A. nidulans* (Δ*thiA*) mono-culture or co-cultivated with *B. subtilis* (WT, Δ*thi*, or Δ*hag*) on minimal medium with/without thiamine grown for 2 days at 30°C. The area of colonies is measured by ImageJ software. Error bar: S.D., n = 3, *** P < 0.001, * P < 0.05. (C) Dispersal of *B. subtilis* (WT or Δ*hag* with ZsGreen) on colonies of *A. nidulans* (Δ*thiA*) without thiamine grown for 2 days at 30°C. Scale bar: 100 μm.

The wild-type *B. subtilis* strain spread to the periphery of the co-cultured fungal D *thiA* colony on the plate without thiamine (Fig. 3C). The *B. subtilis* Δ*thi* cells also dispersed to the periphery of the fungal Δ*thiA* colony even though the fungal Δ*thiA* colony showed a severe growth defect. Because flagella are required for bacterial dispersal on the hyphae, the Δ*hag* cells grew at the center of the fungal colony but did not reach the periphery of the fungal Δ*thiA* colony (Fig. 3C). Notably, the fungal growth defect was hardly recovered by co-culture with the non-motile Δ*hag* strain (Fig. 3B). This was confirmed by other three non-motile mutants (Fig. S6A, B). These results indicated that simultaneous bacterial dispersion to the periphery of the fungal colony and supply of thiamine were required for the normal growth of the *A. nidulans* Δ*thiA* strain.

We measured the amount of thiamine in the supernatant or fungal cell extracts of the co-culture and monoculture by LC-MS-MRM analysis (see methods). The supernatant of *B. subtilis* monoculture contained thiamine 140 ± 2 ng/ml, while thiamine was not detected in that of *B. subtilis* Δ*thi* (Fig. 4A), indicating that *B. subtilis* cells synthesized and secreted thiamine in the medium. The amount of thiamine in the supernatant of co-culture with wild-type *A. nidulans* decreased to 94 ± 5 ng/ml. In contrast, the amount of thiamine in the fungal cell extracts in co-culture with wild-type *B. subtilis* is 134 ± 5 ng/g (wet weight), which was higher than that of fungal monoculture, 95 ± 5 ng/g (wet weight). These support a thiamine transfer from *B. subtilis* to *A. nidulans.*

**Fig. 4.**
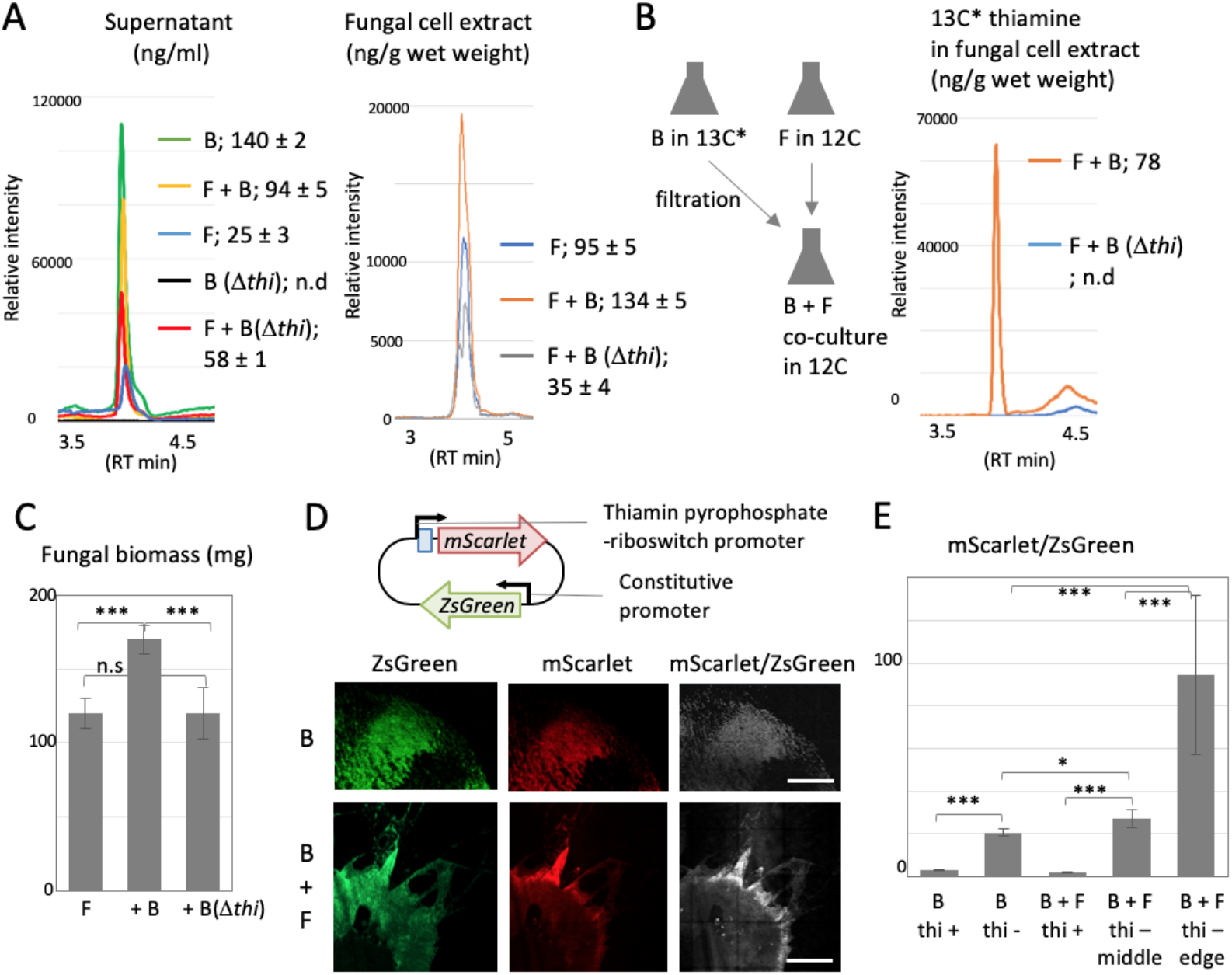
Thiamine transfer analyzed by molecular mass and reporter strain. (A) The amount of thiamine in supernatant and fungal cell extracts in the co-culture and mono-culture of wildtype *A. nidulans* and *B. subtilis* (WT or Δ*thi*) by LC-MS-MRM analysis. B: *B. subtilis,* F: *A. nidulans.* The mean values of peak and S.D. are shown. n = 3. (B) The amount of ^13^C* thiamine by LC-MS-MRM analysis in the fungal cell extracts in the co-culture of *A. nidulans* pre-grown in ^12^C and *B. subtilis* (WT or Δ*thi*) pre-grown in ^13^C*. (C) The fungal biomass in wild-type *A. nidulans* mono-culture and co-culture with *B. subtilis* (WT or Δ*thi*). Error bar: S.D., n = 3, *** P < 0.001. (D) Construct of *B. subtilis* thiamine reporter strain. Colonies of the *B. subtilis* reporter strain mono-culture and co-culture with *A. nidulans* on the minimal medium without thiamine grown for 2 days at 30°C. The images are constructed by 10 x 10 tiling of 500 x 500 μm confocal image. Scale bar: 500 μm. (E) Ratio of signal intensity, mScarlet-1/ZsGreen, in 500 x 500 μm confocal image normalized by ZsGreen intensity. Error bar: S.D., n = 3, *** P < 0.001. * P < 0.05.

The *B. subtilis* wild-type or Δ*thi* cells were labeled by culturing in medium containing stable isotope ^13^C-glucose, while the *A. nidulans* was cultured in medium containing normal glucose. After the 2 day monoculture, the cells were washed and co-cultured. The LC-MS analysis detected ^13^C thiamine in the washed fungal cell extracts in the co-culture with the wildtype *B. subtilis* (Fig. 4B), but not in the co-culture with *B. subtilis* Δ*thi*. These results directly demonstrate that *A. nidulans* cells take the thiamine up from *B. subtilis.* Indeed, the fungal biomass in the co-culture was 40 % higher than that in the fungal monoculture (Fig. 4C), consisting with Fig. 2A. In contrast, the amount of thiamine in fungal cell extracts and the fungal biomass did not increase in the co-culture with *B. subtilis* Δ*thi*. These indicate that the supply of thiamine from *B. subtilis* promotes the fungal growth.

We constructed a *B. subtilis* thiamine reporter strain, expressing ZsGreen, under the constitutive promoter, and mScarlet-1, under the control of thiamin pyrophosphate (TPP)-riboswitch, whose expression is activated in the thiamine-depleted condition (Fig. 4D) (21). The mScarlet-1 was not expressed in the bacterial colony grown with thiamine but induced without thiamine (Fig. 4D, and Fig. S7A). In co-culture with *A. nidulans* as well, the mScarlet-1 was not expressed with thiamine but induced without thiamine. Notably, the induction was significantly higher at the edge of colony than in the middle (Fig. 4D, E, and Fig. S7A, B). These indicate that *B. subtilis* cells produce more thiamine at the colony edge because *A. nidulans* takes thiamine up at the growing hyphal tips.

Taken together with our results, bacterial benefit is the bacterial cells moving faster along hyphae and the hyphae delivering the bacteria farther, while the fungal benefit is delivery of thiamine to hyphal tips by bacterial cells and resultant promotion of fungal growth.

The LC-MS-MRM analysis indicated the amount of thiamine in the fungal cell extracts in co-culture with *B. subtilis* Δ*thi* was lower than that of fungal monoculture (Fig. 4A). CFU of *B. subtilis* in the co-culture of *B. subtilis* Δ*thi* and *A. nidulans* wild-type was higher than that in monoculture of *B. subtilis* Δ*thi*, while that in co-culture *B. subtilis* Δ*thi* and *A. nidulans* Δ*thiA* was comparable with monoculture of *B. subtilis* Δ*thi* (Fig. S6D). These results indicate bidirectional thiamine transfer between *B. subtilis* and *A. nidulans*.

### Ecological relevance of mutualistic interaction

To evaluate the ability of fungi to disperse bacteria in nature, we observed bacterial movement along the hyphae in water-unsaturated conditions, on glass surfaces and in soil (Fig. S8 and Movie S11-13). In addition, we designed a soil-sandwich experiment as follows. A square section of agar with co-cultured *A. nidulans* and *B. subtilis* (right) and another new section of agar (left) were placed a few mm apart; the separation between the two sections was filled with soil particles (Fig. 5A). The fungal hypha protruding from the right agar, continued to extend into the soil particles, and eventually reached the left agar slab (Fig. 5A and Movie S14). Migration of *B. subtilis* cells (green) followed the mycelium extension, through the soil particles, and to the left agar slab. In the absence of the fungus or soil particles, no bacteria migrated beyond the gap between agar sections (Fig. S9 and Movie S15, 16). These indicate that hyphal growth towards favorable nutrient conditions on dry solid substrates enables bacteria to move along the hyphae and explore previously inaccessible spatial niches in nature.

**Fig. 5.**
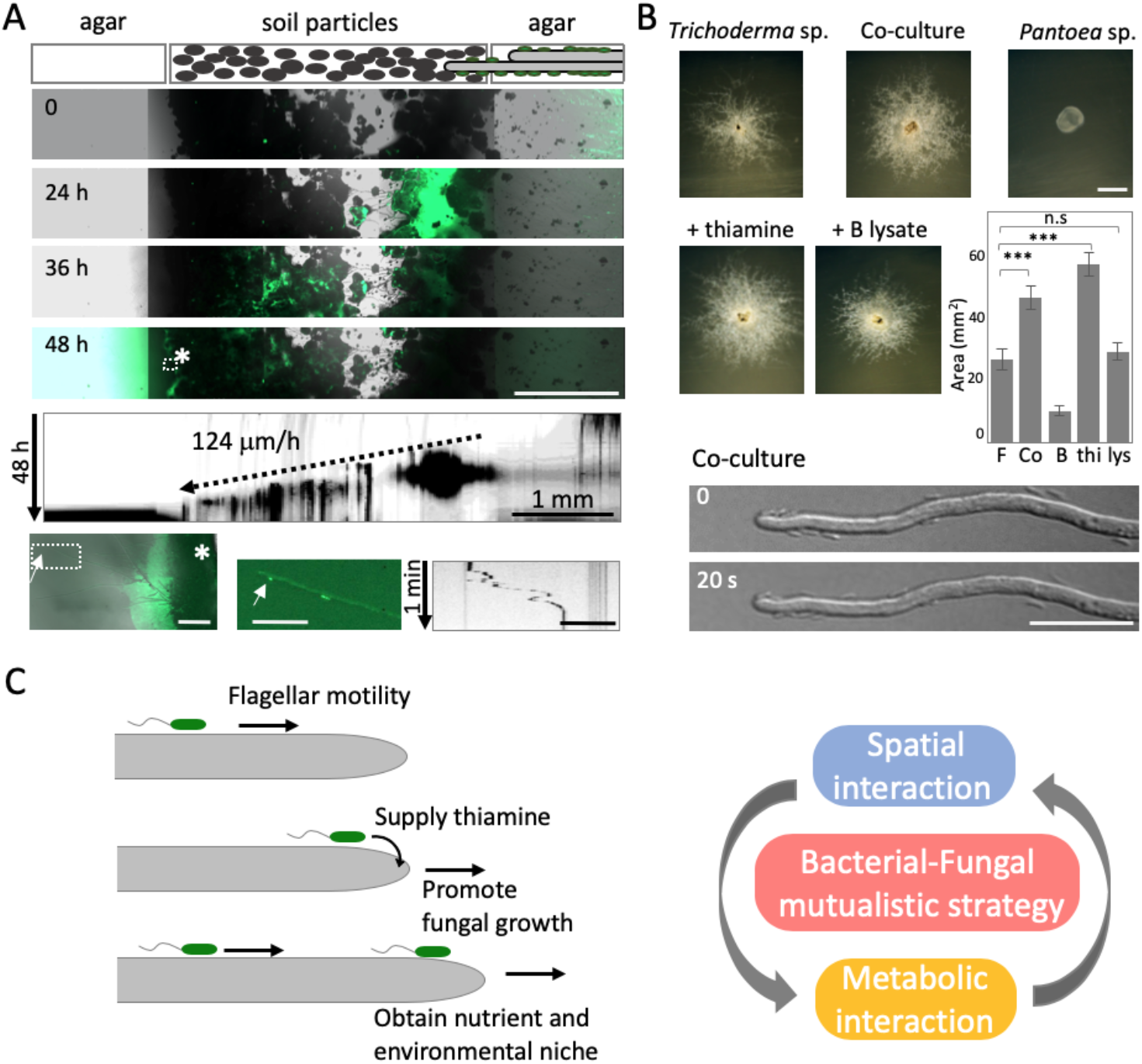
Mutualistic growth strategy by spatial and metabolic interactions. (A) Time-lapse bacterial dispersal (green) on growing hyphae in soil particles sandwiched between two agar pieces at 0, 24, 36, and 48 h. Scale bar: 1 mm. Kymograph of bacterial migration from Movie S14 (Vertical arrow: 48 h, Scale bar: 1 mm). Asterisk indicates an expanded image of bacterial colony (green) and mycelium at the left agar after 48 h (left bottom). Scale bar: 100 μm. Arrows indicate bacterial movement along hyphae at the left agar after 48 h (middle bottom). Scale bar: 50 μm. Kymograph of the bacterial movement (right bottom). Vertical arrow: 1 min, Scale bar: 50 μm. (B) Colonies *Trichoderma* sp. and *Pantoea* sp. mono-culture and co-culture. The bacterial cell lysate is prepared by sonication. Scale bar: 2 mm. The area of colonies are measured by ImageJ software. Error bar: S.D., n = 3, *** P < 0.001. Expanded image of the bacterial cells move along the hyphae and reach the tip. Scale bar: 20 μm. (C) Mutualistic growth strategy that the bacterial cells move faster along fungal highway and disperse farther on fungal growth, while bacterial cells supply thiamine to hyphal tips and promote the fungal growth.

To confirm the ecological relevance, we screened a bacterial-fungal complex, where bacteria moved along fungal hypha, from natural soil and co-isolated *Trichoderma* sp. (*harzianum* and *neotropicale*; 100% identity of 255 bp ITS) and *Pantoea* sp (98.4% similarity to the full length of 16S rRNA gene in *P. rodasii*). The fungus co-cultured with bacteria grew better than the fungus mono-culture (Fig. 5B). In the co-culture, the bacteria cells moved along the hyphae and reached the tips of hyphae (Movie S17). The fungal growth was promoted with the addition of thiamine to the whole medium, but not with spot inoculation of bacterial cell lysates, which mimic a non-motile mutant. This example supports the ecological relevance of similar mechanism observed in the co-cultured *A. nidulans* and *B. subtilis*.

## Discussion

Here we displayed bacterial motility along fungal hyphae and bacterial dispersal on mycelial extension. The phenomena were observed on agar media, glass and in the soil, which are water unsaturated conditions. It is likely that water surrounds and covers the hyphae due to surface tension, thus suppling space around hyphae for bacterial cells to swim by their flagella. Mycelium networks extend in natural environment, especially throughout soil, which could function as roads and bases for bacteria to migrate and proliferate. This is consistent with a recent study showing that fungal networks shape the dispersal of bacteria in the cheese rind microbiota (22).

While previous works have demonstrated the bacterial transportation via fungal highway (13,14, 23), several metabolic interactions have been analyzed between fungi and bacteria (6–8, 24). Our transcriptomic, genetic, molecular mass and imaging analyses propose a mutualistic growth mechanism that the bacterial cells move faster along the fungal highway and disperse farther in the niche, concomitantly the bacterial cells reach the mycelial edge and supply thiamine to the growing hyphae, resulting in a promotion of hyphal growth (Fig. 5C). The simultaneous spatial and metabolic interactions, that are the bacterial dispersal on fungal highway and share of thiamine, establish a mutualism that facilitates the communicating fungal and bacterial species to obtain environmental niche and nutrient respectively (Fig. 5C). Although the bacterial-fungal combination we tested is an artificial condition, the example of co-isolated bacterial-fungal species from nature supports the ecological relevance of the mutualism through fungal highway and share of thiamine.

It has been recently reported that *B. subtilis* supply thiamine to a thiamine-auxotrophic fungus, which is an endophytic fungus colonizing to roots of wide-range plants and has lost genes related to thiamine synthesis (25). The different aspect of our finding is that *A. nidulans* can synthesize thiamine on their own but utilize thiamine from *B. subtilis*. Thiamine is an essential co-factor for central carbon metabolism in all living organisms and is synthesized by bacteria, fungi, and plants (26). Since thiamine often limits the growth of these organisms, they have evolved numerous strategies to obtain thiamine from the natural environment (27). Thiamine riboswitches are one of the strategies used to tightly regulate thiamine synthesis and uptake (28). Since we find the bacteria secrete thiamine extracellularly, the neighboring non-auxotrophic bacteria and fungi, like *A. nidulans,* in nature could utilize the thiamine by uptake to save the cost rather than by synthesis. Thiamine and riboswitch have the potential to be used widely and universally stimulating symbiosis among microbes and even inter-kingdom interactions in nature. Other vitamins besides thiamine, which are essential for growth but sufficient in small amounts, are likely the seeds of commensal and mutualistic interactions of microbes in nature (29–32).

The affinity of fungal-bacterial interactions is selective depending on the combination of species (unpublished data). Besides natural auxotrophy, secondary metabolites are also involved in microbial communication for selective interaction. Especially soil-dwelling bacteria and fungi produce a wide range of secondary metabolites, which function as communication signals among microorganisms to compete and interact with others (33). Some secondary metabolite genes are upregulated in *A. nidulans* co-cultured with *B. subtilis.* The combined analysis of natural auxotrophy and secondary metabolites in co-culture of bacteria and fungi will provide hints to understand selective microbial communication. In addition, live imaging of bacterial-fungal co-culture represents an efficient approach to bioassays that screen for affinities between bacteria and fungi. Recent studies indicate coordinated interactions between fungi and bacteria in various situations, such as promotion of plants growth, fermentation, biomass degradation, plant pathogenesis, and human pathogenesis (5, 6, 22, 34). Although most studies reveal the metabolic interactions, besides them, imaging the localization and functional distribution of microbes is increasing in importance.

## Materials and Methods

### Strains and media

A list of *A. nidulans* and *B. subtilis* strains used in this study is given in Table S4. Supplemented minimal medium for *A. nidulans* and standard strain construction procedures were described previously (35). If necessary, thiamine was complemented at 10 μM. Surface soil sample was collected at the depth of (0-5 cm) in secondary forest in University of Tsukuba, Japan. The soil sample was gently sieved by 250 μm stainless steel mesh in a field moist condition.

### Strain construction

*B. subtilis* Δ*thi* and Δ*hag* strains were constructed as follows. 500 bp of flanking regions were amplified by PCR using primer sets tenA-5/-N3 and tenA-C5/-3, and hag-N5/-N3 and hag-C5/-3, respectively (Table S5). Antibiotic resistance genes *Cat^R^* and *Spc^R^* were amplified using the primer sets cat-Fw/-Rv and spc-Fw/-Rv. The three fragments were ligated by overlap PCR. The DNA fragments were then transformed into *B. subtilis* 168 to construct Δ*thi* [*Cat^R^*] and Δ*hag* [*Spc^R^*] strains. The deletion of target genes was confirmed by PCR and sequencing. *B. subtilis* expressing green or red fluorescent protein maintains the plasmid, pHY300-Pveg-ZsGreen-term [*Tet^R^*] or pHY300-Pveg-mCherry-term [*Tet^R^*](36). *B. subtilis* thiamine reporter strain (thi-rep) maintains the plasmid, pHY300MK, expressing ZsGreen under the constitutive Pveg and mScarlet-1 under the promoter of *tenA-operon* 300 bp amplified using the primer set tenA-5/-mSca-Rv.

### Microscopy

A confocal laser scanning microscope (CLSM) LSM880 (Carl Zeiss, Jena, Germany) equipped with a 63×/0.9 numerical aperture Plan-Apochromat objective and a 40x/0.75 numerical aperture IR Achroplan W water immersion objectives (Carl Zeiss), were used to acquire confocal microscopic images. Bacteria and/or fungi were irradiated with 488 and 633-nm lasers to detect the Green fluorescent protein (ZsGreen) and reflected light, respectively. Acquired confocal images were analyzed using ZEN Software (Version 3.5, Carl Zeiss) and ImageJ software. Another CLSM TCS SP8 Tandem scanner 8kHz (Leica, Mannheim, Germany) equipped with HC PL APO 20x/0.75 IMM CORR CS2 objective lens was used to acquire hi-speed confocal dual-color microscopic images. Bacteria and fungi were irradiated with 488 and 522-nm lasers to detect the GFP and mCherry, respectively. Epifluorescent inverted microscopy: cells were observed with an Axio Observer Z1 (Carl Zeiss) microscope equipped with a Plan-Apochromat 63×1.4 Oil or 10 or 20 times objective lens, an AxioCam 506 monochrome camera and Colibri.2 LED light (Carl Zeiss). Temperature of the stage was kept at 30°C by a thermo-plate (TOKAI HIT, Japan). Zoom microscopy; Plates were observed by AXIO Zoom V16 and HXP 200C illuminator (Carl Zeiss). Images were collected and analyzed using the Zen system (Carl Zeiss) and ImageJ software.

### RNA-seq analysis

Total RNA was isolated from fungal and bacterial cells that were cocultured for 8 h in 6-well plates. For bacterial RNA isolation, cells were disrupted using glassbeads. For fungal RNA isolation, cells were frozen and homogenized using mortar and pestle, and then the total RNA was extracted using an RNA isolation kit (RNeasy Mini Kit, Qiagen). Ribosomal RNA was depleted using Ribo-ZERO magnetic kit (Epicentre). The transcripts were fragmented and used as templates to generate strand-specific cDNA libraries by TruSeq Stranded Total RNA LT Sample Prep kit (Illumina). Each sample was sequenced using 100-bp paired-end reads on an Illumina HiSeq 2500 instrument. Macrogen Inc. supported library preparation, sequencing and partial data analysis. The reads were mapped to reference genomes of *B. subtilis* NCIB3610 (CP020102.1) or *A. nidulans* TN02A3 (GCA_000149205.1) with Bowtie 2 aligner. Read count per gene was extracted from known gene annotations with HTSeq program. After log2 transformation of RPKM+1 and quantile normalization, differentially expressed genes were selected on conditions of log2 > 2 in expression level.

### Extraction of thiamine

Monocultures of the *A. nidulans* or *B. subtilis* strains and co-cultures were grown in the minimal medium 200 ml with 120 rpm shaking at 30°C for 3 days prior extraction. Thiamine extraction from the cells: the cells were sieved using Mira cloth and freeze dried. Then they were frozen with liquid nitrogen and crushed. The resultant pellet was dissolved in 5ml 0.1M HCl and heated at 100°C for 15 min. It was filtered and allowed to cool. The filtrate was then freeze dried. Finally, freeze-dried pellet was dissolved in 200 μl of preprepared solvent (10 mM ammonium formate + 1 % Methanol + 0.1 μl formic acid). Thiamine extraction from the supernatants: the supernatant was collected after centrifuge 8,000 rpm for 5 min and freeze dried. The resultant pellet was dissolved in the pre-prepared solvent.

### ^13^C labeling

*B. subtilis* wild-type or Δ*thi* cells were cultured in the minimal medium containing [U-13C6, 99%] labeled D-glucose (1%) [Cambridge Isotope Laboratories, Inc.] in 200 ml with 120 rpm shaking at 30°C for 2 days. Simultaneously *A. nidulans* were cultured in minimal medium containing normal D-glucose in 200ml for 2 days. The *B. subtilis* cells were collected by centrifuge, while the *A. nidulans* cells were collected using Mira cloth and washed. Then they were co-cultured in minimal medium containing D-glucose 200ml for 2 days. The fungal cells were sieved through Mira cloth and washed thoroughly with milliQ water to remove any bacterial cells attached to the surface. Extraction of labeled thiamine from the cell extract followed the same protocol as of extraction of thiamine described above.

### LC-MS-MRM analysis

Resultant pellets that were dissolved in the pre-prepared solvent were analyzed by LC-MS (LCMS 8030, Shimadzu, Kyoto, Japan) equipped with a 250 x 3.0 mm COMOSIL HILIC Packed Column (particle size 5 μm; Nacalai Tesque, INC, Japan). The initial mobile phase was with a ratio of solvent A: solvent B (solvent A-Acetontrile:10 mM Ammonium acetate in Water (9:1); solvent B-100 % Acetonitrile), increased to 100 % and maintained at that ratio for 7 min. UV/Vis spectra were monitored by SPD-M30A (Shimadzu). The mass spectrometer was operated in Multiple Reaction Monitoring (MRM) mode for quantitative analysis of thiamine in the corresponding samples. Mass spectra were acquired with the following conditions: capillary voltage 4.5kV; detection range m/z 122 for normal thiamine and 128 (Precursor m/z 277) for ^13^C labeled thiamine; desolvation line 250 °C; heat block 400 °C; nebulizer gas 3 l/min; drying gas 15 l/min. Calibration curves were obtained using the LabSolution software (v5.91 Shimadzu Corporation, Japan).

### Fungal biomass

Monoculture of *A. nidulans* and co-cultures with *B. subtilis* wild-type and Δ*thi* strains were grown in the minimal medium 100 ml with 120 rpm shaking at 30°C for 3 days. The cultures were then filtered using Mira cloth and the pellet was washed with milliQ water thoroughly (to remove the bacteria in the coculture filtrates). The resultant pellets were then freeze dried (SCANVAC COOLSAFE, LaboGene, Allerød). The weight of the dried pellets was measured several times in between freeze drying until the weight was constant.

### Bacterial genomic DNA extraction

Monoculture and co-cultures were grown on cellophane film on the minimal medium agar (point inoculation of OD_600_ = 0.01). The plates were incubated at 30 °C for 3 days. The cellophane films were washed with sterilized milliQ carefully and thoroughly to collect the bacterial cells. The collected samples were subjected to bacterial genomic DNA extraction protocol using Wizard^®^ Genomic DNA purification kit. The purified genomic DNA was quantified using nanodrop (Thermoscientific nanodrop 2000, ThermoFisher, USA).

### MATLAB

ImageJ was used to generate a list of cell centroid positions (*x*_i_, *y*_i_), where *i* is the frame index number. MATLAB (Natick, MA) was used to calculate the instantaneous speed of each cell and then speed heat maps were generated. These heat maps were plotted as function of instantaneous position (see Fig. 1C), while speed was plotted as a function of time. The underlying motion was extracted using a moving window average of 5 consecutive values to smooth the data and then fitted using a sinusoid.

### Statistical analysis

Student t-tests were used to evaluate the mean difference between two sets.

## Data availability

All data generated or analyzed during this study were included in the manuscript and Supplementary Information. The original data generated during and/or analyzed during this study are available from the corresponding author on reasonable request.

## Acknowledgments

We appreciate the experimental support of Hiroko Kato and Takamitsu Soma at University of Tsukuba. This work is supported by Japan Society for the Promotion of Science (JSPS) KAKENHI grant number 18K05545 and 18K15143, a grant from the Institute for Fermentation, Osaka (IFO), Noda Institute for Scientific Research Grant and Japan Science and Technology Agency (JST) ERATO grant number JPMJER1502.

## Authors contributions

N.O., N.N., and N.T. designed the research project. G.A., M.K., G.K., and N.T. performed microscopy experiments and analysis of data. K.T., and A.U. developed the MATLAB analysis. G.A., A. N. and S.M. performed LC-MS and analyzed data. N.O., M.K., and D.H. performed RNA-seq and analysis of data. N.O., N.T. and N.T. wrote the manuscript with inputs from other coauthors.

## Competing interests

The authors declare that they have no competing interests.

## Notes

### Competing Interest Statement

The authors have declared no competing interest.

## References

1. Romine MF, Rodionov DA, Maezato Y, Osterman AL, Nelson WC (2017) Underlying mechanisms for syntrophic metabolism of essential enzyme cofactors in microbial communities. ISME J. 11: 1434–1446.

2. Zengler K, Zaramela LS (2018) The social network of microorganisms - how auxotrophies shape complex communities. Nat. Rev. Microbiol. 16: 383–390.

3. Osono T, Ono Y, Takeda H (2003) Fungal ingrowth on forest floor and decomposing needle litter of Chamaecyparis obtusa in relation to resource availability and moisture condition. Soil. Biol. Biochem. 35: 1423–1431.

4. Fierer N (2017) Embracing the unknown: disentangling the complexities of the soil microbiome. Nat. Rev. Microbiol. 15: 579–590.

5. Nazir R, Warmink JA, Boersma H, van Elsas JD, (2010) Mechanisms that promote bacterial fitness in fungal-affected soil microhabitats. FEMS. Microbiol. Ecol. 71: 169–185.

6. Frey-Klett P, et al. (2011) Bacterial-fungal interactions: hyphens between agricultural, clinical, environmental, and food microbiologists. Microbiol. Mol. Biol. Rev. 75: 583–609.

7. Benoit I, et al. (2015). *Bacillus subtilis* attachment to *Aspergillus niger* hyphae results in mutually altered metabolism. Environ Microbiol 17: 2099–113

8. Worrich A, et al. (2017) Mycelium-mediated transfer of water and nutrients stimulates bacterial activity in dry and oligotrophic environments. Nat. Commun. 8: 15472.

9. Nutzmann HW, et al. (2011) Bacteria-induced natural product formation in the fungus *Aspergillus nidulans* requires Saga/Ada-mediated histone acetylation. Proc. Natl. Acad. Sci. U.S.A. 108: 14282–14287.

10. Riquelme M, et al. (2018) Fungal morphogenesis, from the polarized growth of hyphae to complex reproduction and infection structures. Microbiol. Mol. Biol. Rev. 82: 1–47.

11. Kearns DB (2010) A field guide to bacterial swarming motility. Nat. Rev. Microbiol. 8: 634–644.

12. Harshey RM, (2003) Bacterial motility on a surface: many ways to a common goal. Annu. Rev. Microbiol. 57: 249–273.

13. Kohlmeier S, et al. (2005) Taking the fungal highway: mobilization of pollutantdegrading bacteria by fungi. Environ. Sci. Technol. 39: 4640–4646.

14. Pion M, et al. (2013) Gains of bacterial flagellar motility in a fungal world. Appl. Environ. Microbiol. 79: 6862–6867.

15. Takeshita N, et al. (2017) Pulses of Ca^(2+)^ coordinate actin assembly and exocytosis for stepwise cell extension. Proc. Natl. Acad. Sci. U.S.A. 114: 5701–5706.

16. Zhou L, et al. (2018) Superresolution and pulse-chase imaging reveal the role of vesicle transport in polar growth of fungal cells. Sci. Adv. 4: e1701798.

17. Cazorla FM, et al. (2007) Isolation and characterization of antagonistic *Bacillus subtilis* strains from the avocado rhizoplane displaying biocontrol activity. J. Appl. Microbiol. 103: 1950–1959.

18. Matsuura S, Shioi JI, Imae Y (1977) Motility in *Bacillus subtilis* driven by an artificial protonmotive force. FEBS. Lett. 82: 187–190.

19. Kearns DB, Losick R (2003) Swarming motility in undomesticated *Bacillus subtilis*. Mol. Microbiol. 49: 581–590.

20. Shimizu M, et al. (2016) Thiamine synthesis regulates the fermentation mechanisms in the fungus *Aspergillus nidulans*. Biosci. Biotechnol. Biochem. 80: 1768–1775.

21. Mironov AS, et al. (2002) Sensing small molecules by nascent RNA: a mechanism to control transcription in bacteria. Cell 111: 747–756.

22. Zhang Y, Kastman EK, Guasto JS, Wolfe BE (2018) Fungal networks shape dynamics of bacterial dispersal and community assembly in cheese rind microbiomes. Nat. Commun. 9: 336.

23. Ingham CJ, Kalisman O, Finkelshtein A, Ben-Jacob E (2011) Mutually facilitated dispersal between the nonmotile fungus *Aspergillus fumigatus* and the swarming bacterium *Paenibacillus vortex*. Proc Natl Acad Sci USA 108(49):19731–36

24. Deveau A, Brulé C, Palin B, Champmartin D, Rubini P, Garbaye J, Sarniguet A, Frey-Klett P. (2010). Role of fungal trehalsoe and bacterial thiamine in the improved survival and growth of the ectomycorrhizal fungus *Laccaria bicolor* S238N and the helper bacterium *Pseudomonas fluorescens* BBc6R8. Environ Microbiol Rep 2(4):560–8

25. Jiang X, et al. (2018) Impact of spatial organization on a novel auxotrophic interaction among soil microbes. ISME J. 12: 1443–1456.

26. Jurgenson CT, Begley TP, Ealick SE (2009) The structural and biochemical foundations of thiamin biosynthesis. Annu. Rev. Biochem. 78: 569–603.

27. Kraft CE, Angert ER (2017) Competition for vitamin B 1(thiamin) structures numerous ecological interactions. Q. Rev. Biol. 92: 151–68.

28. Cressina E, et al. (2011) Identification of novel ligands for thiamine pyrophosphate (TPP) riboswitches. Biochem. Soc. Trans. 39: 652–657.

29. Klein CC, et al. (2013) Biosynthesis of vitamins and cofactors in bacterium-harbouring trypanosomatids depends on the symbiotic association as revealed by genomic analyses. PLoS One. 8: e79786.

30. Magnúsdóttir S, Ravcheev D, de Crécy-Lagard V, Thiele I. (2015) Systematic genome assessment of B-vitamin biosynthesis suggests co-operation among gut microbes. Front Genet. 6:148.

31. Palacios OA, et al. (2016) Tryptophan, thiamine and indole-3-acetic acid exchange between Chlorella sorokiniana and the plant growth-promoting bacterium Azospirillum brasilense. FEMS Microbiol Ecol. 92: fiw077.

32. Sokolovskaya OM, Shelton AN, Taga ME. Sharing vitamins: Cobamides unveil microbial interactions. Science. 2020;369

33. Macheleidt J, et al. (2016) Regulation and Role of Fungal Secondary Metabolites. Annu. Rev. Genet. 50: 371–392.

34. Barkal LJ, et al. (2017) Microbial volatile communication in human organotypic lung models. Nat. Commun. 8: 1770.

35. Takeshita N, et al. (2013) The cell-end marker TeaA and the microtubule polymerase AlpA contribute to microtubule guidance at the hyphal tip cortex of *Aspergillus nidulans* to provide polarity maintenance. J. Cell Sci. 126: 5400–11.

36. Toyofuku M, et al. (2017) Prophage-triggered membrane vesicle formation through peptidoglycan damage in *Bacillus subtilis*. Nat. Commun. 8: 481.

