## Supplement figures, tables, movies for "Fungal mycelia and bacterial thiamine establish a mutualistic growth mechanism": Supplementary.pdf

*Correspondance to:

Nozomu Obana,

Norio Takeshita,

[#]Equally contribution

[†]Current position, MiCS and Transborder Medical Research Center, Faculty of Medicine, University of Tsukuba

**Fig. S1-9**

**Table S1-5**

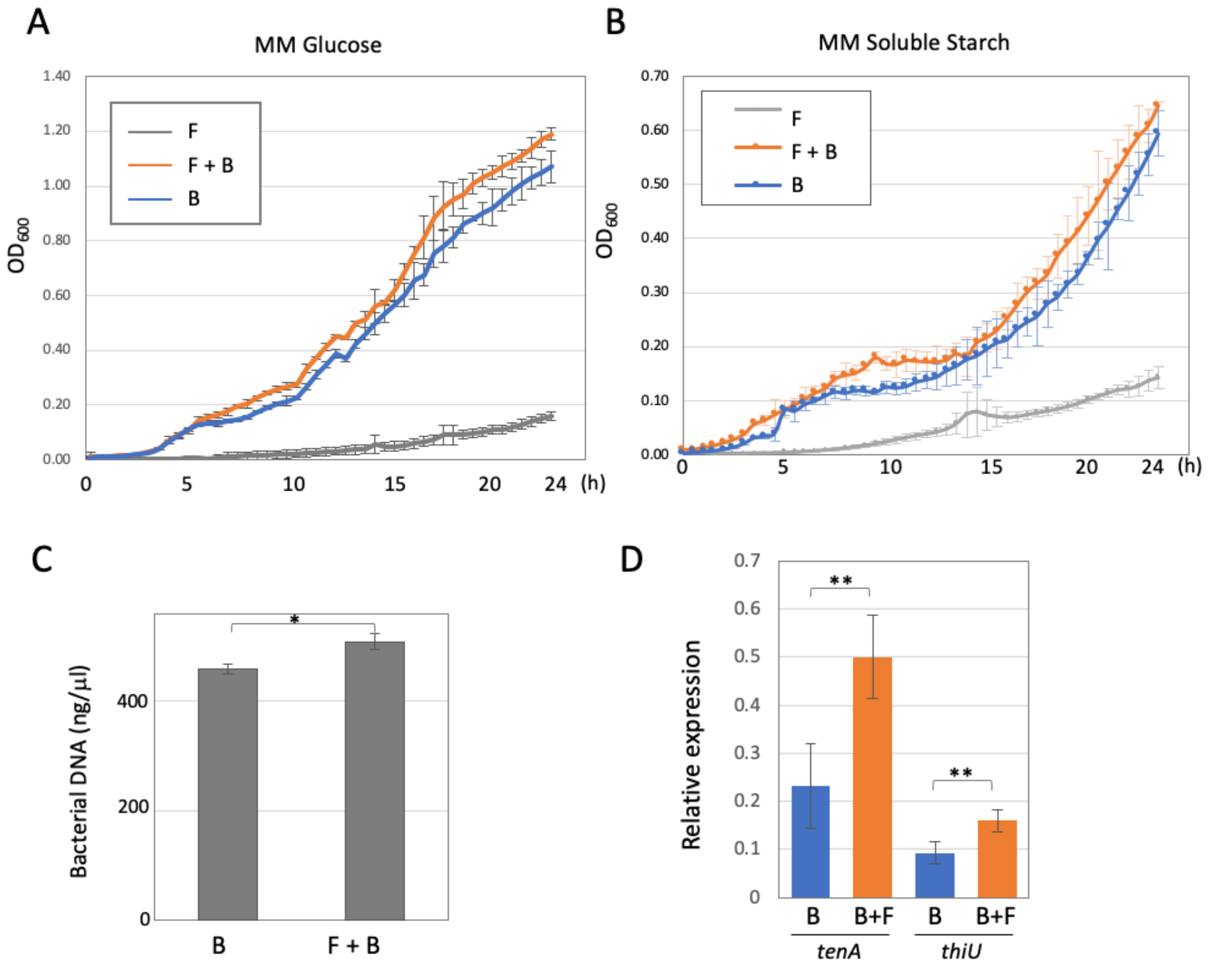

**Fig. S1. Effects of co-culture with the fungus on the bacterial growth.** (A, B) The spores of *A. nidulans* were incubated in 200 µl minimal medium with glucose (A) or amylose (B) for 7 hours until they germinated, then pre-cultured *B. subtilis* were inoculated at a final concentration of $OD_{600}$=0.01. Bacterial growth rates measured in 96-well titer plate and shown by $OD_{600}$ every 30 min for 24 h. Error bar: S.D., n = 3. F: fungus, B: bacteria. (C) Bacterial genomic DNA was extracted from bacterial mono-culture or co-culture with the fungus on the minimal medium agar plates incubated at 30 °C for 3 days. Error bar: S.D., n = 3. * P≤ 0.05. (D) The relative expressions of *tenA* and *thiU* in the bacterial mono-culture or co-culture with the fungus measured by qRT-PCR. The expression of *sigA* is used as a standard. Error bar: S.D., n = 3. ** P≤ 0.01.

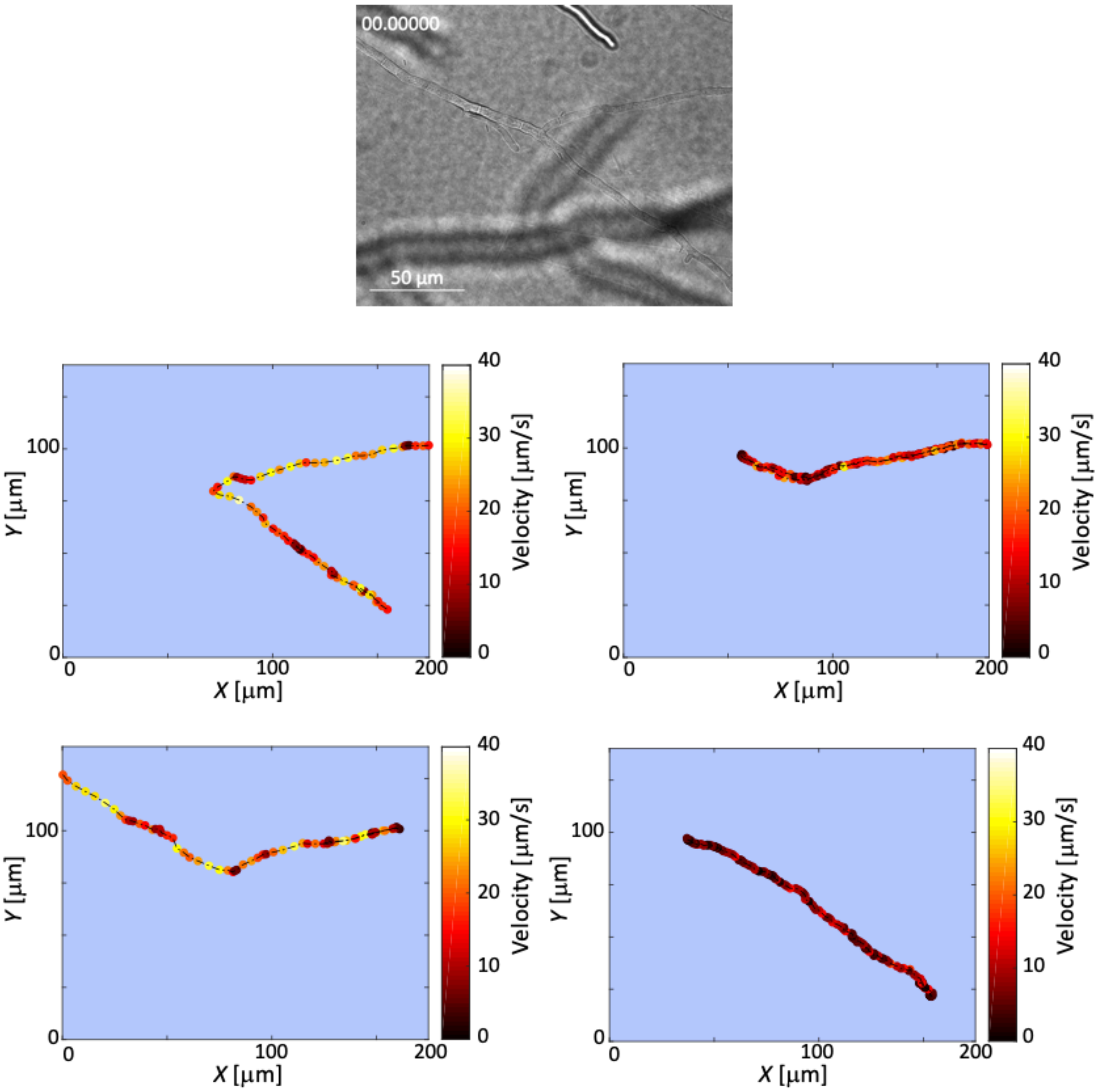

**Fig. S2. Oscillations in *B. subtilis* movement along hyphae.** Heat-maps of *B. subtilis* instantaneous velocity analyzed by track the position of each cell moving along hyphae from Movie S1. We tracked the positions of each cell moving along hyphae and from the instantaneous position of the center of mass of a cell. We generated heat-maps of the instantaneous velocity, where darker colors represent slower speeds and lighter colors represent higher velocities, respectively. The heat-maps indicate that there is a weak oscillation in the instantaneous velocity over time (Fig. 1C). The reason for the oscillations that we see in the instantaneous velocity of bacteria moving along hyphae may be the result of stick-slip motion mediated by the flagella. Due to the low water content and narrow gaps between hyphae and agar, the cells may momentarily become wedged in tight gaps; however, the flagellar motors may exert sufficient force to free the stuck cells.

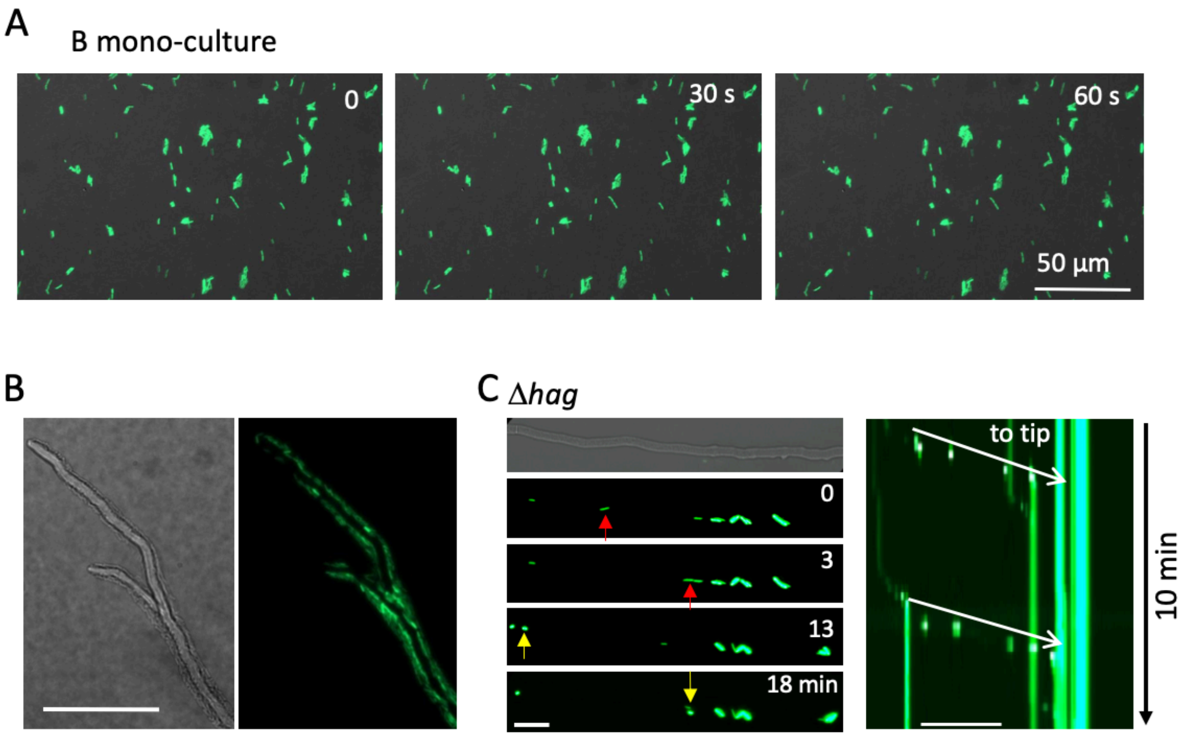

**Fig. S3.** ***B. subtilis* movement along hyphae by flagella.** (A) Time-lapse images of *B. subtilis* (green) mono-culture for 60 s on the minimum agar media. Scale bar: 50 µm. (B) *A. nidulans* hyphae (DIC) surrounded by moving *B. subtilis* (green) from Movie S4. (C) Image sequence of *B. subtilis* flagella mutant (Δ*hag*) flow (arrows) along *A. nidulans* hyphae. Kymograph of the Δ*hag* along the hyphae to the tip (white arrows) from Movie S5. Total 10 min. Scale bar: 50 µm.

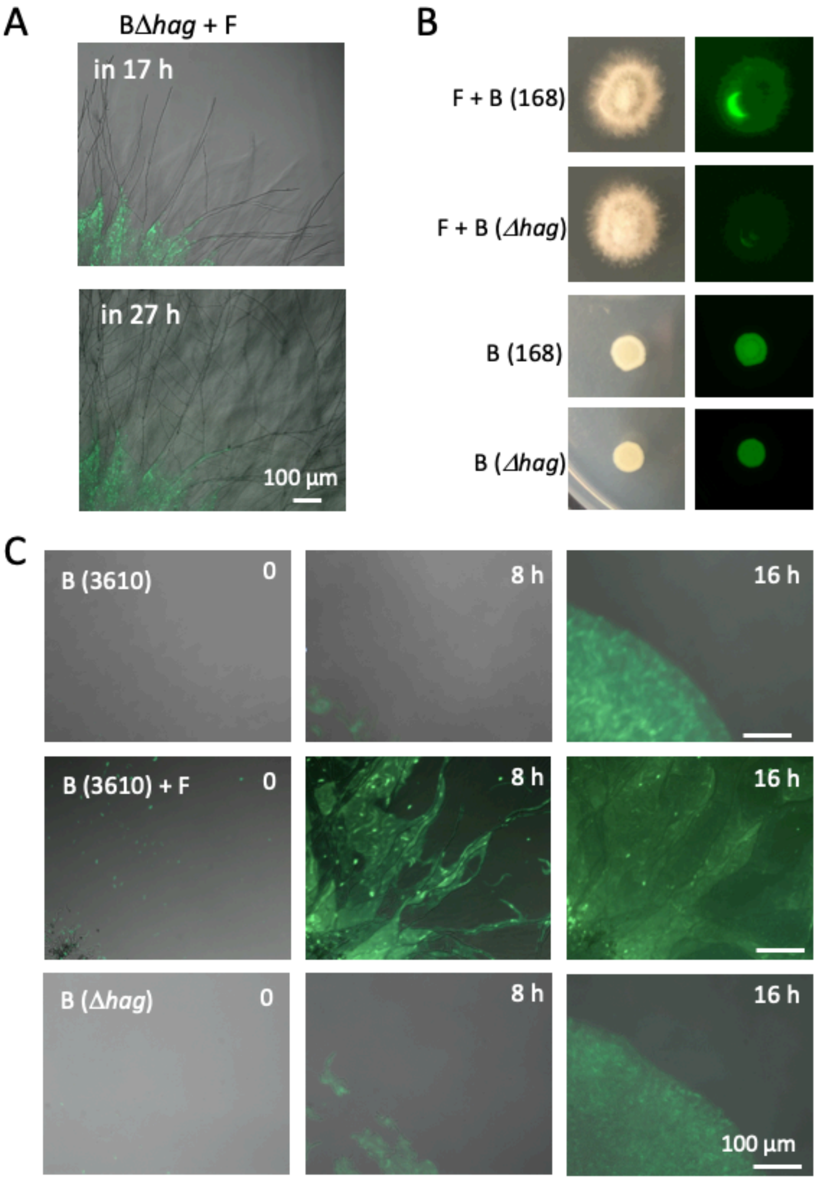

**Fig. S4. Bacteria dispersal on fungal colony by flagella.** (A) Time-lapse proliferation of *B. subtilis* Δ*hag* (green) and *A. nidulans* (DIC) after 17 h of co-culture. Scale bar: 100 μm. (B) Bright field (left) and green fluorescent (right) images of colonies of *B. subtilis* (168; WT or Δ*hag*) mono- or co-culture with *A. nidulans*. Aerial growing hypha at the middle of colony disturb to detect the fluorescent signals in the co-culture. (C) Time-lapse proliferation of *B. subtilis* (3610; WT) mono- or co-culture with *A. nidulans*, and *B. subtilis* (Δ*hag*) mono-culture. Scale bar: 100 μm.

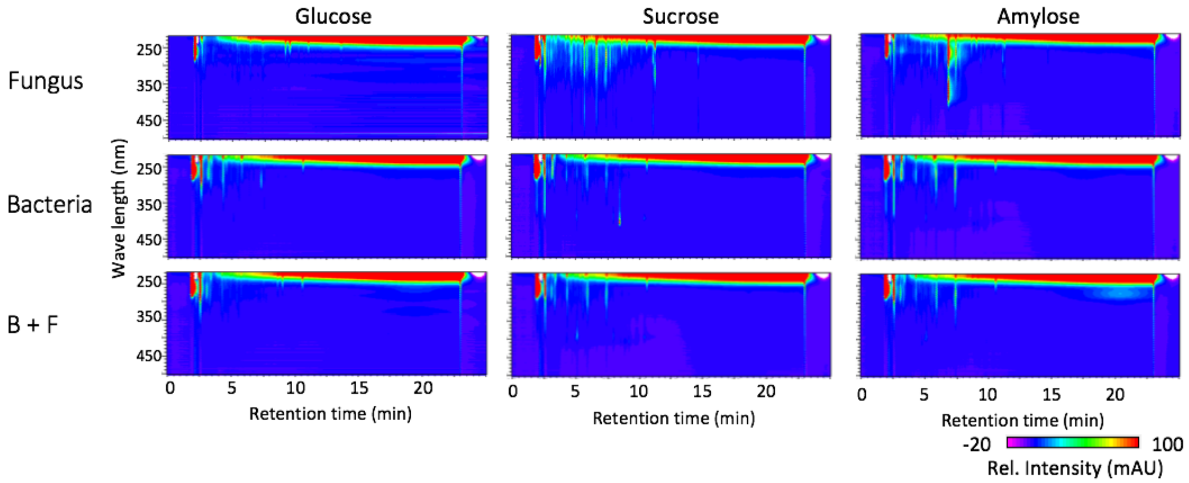

**Fig. S5. Extracellular hydrophobic metabolites in mono- or co-culture with different carbon source analyzed by LC-PDA-ESI/MS.** Each microorganism was cultured in minimum medium with indicated carbon source for 5 days. Extracellular hydrophobic metabolites in culture supernatant were extracted with acidified ethyl acetate and analyzed by LC-PDA-ESI/MS. Contour maps were constructed from the absorption intensity obtained by LC-PDA analysis. In each mono-culture with different carbon source, we see little change in the EHM profiles of *B. subtilis.* On the other hand, the profiles of *A. nidulans* are affected by the different carbon sources, which consistent with the previous reports about the regulation of secondary metabolism. EHM in co-culture are similar to those in *B. subtilis* mono-culture regardless of carbon source. The fungal and bacterial cells co-cultured as described above for 5 days in 6-well plates. Supernatant of mono- or co-culture was collected by centrifugation at 15,000 x g for 10 min, and acidified by 1/100 volume of 2 M HCl. Equal volume of ethyl acetate was added to the acidified supernatant, stirred for 1 h and centrifuged at 1,000 x g for 10 min. The ethyl acetate fraction was collected to other tubes, and lyophilized. Resulting pellet was dissolved in 95% methanol and analyzed by LC-PDA-ESI/MS (LCMS-8030; Shimadzu, Kyoto, Japan) equipped with a 150 × 4.6-mm Purospher Star RP-18 column (particle size, 5 μm; Millipore-Merck, Billerica, MA, USA). The initial mobile phase was solvent A: solvent B = 98:2 (solvent A, 0.1% formic acid; solvent B, acetonitrile), increased to 80% for x min and maintained at that ratio for another 5 min. UV/Vis spectra was monitored by SPD-M30A (Shimadzu). Mass spectra were acquired in the positive mode of LCMS-8030 with the following conditions: capillary voltage, 4.5 kV; detection range, m/z 50–600; desolvation line, 250°C; heat block, 400°C; nebulizer gas, 3 L/min; drying gas, 15 L/min.

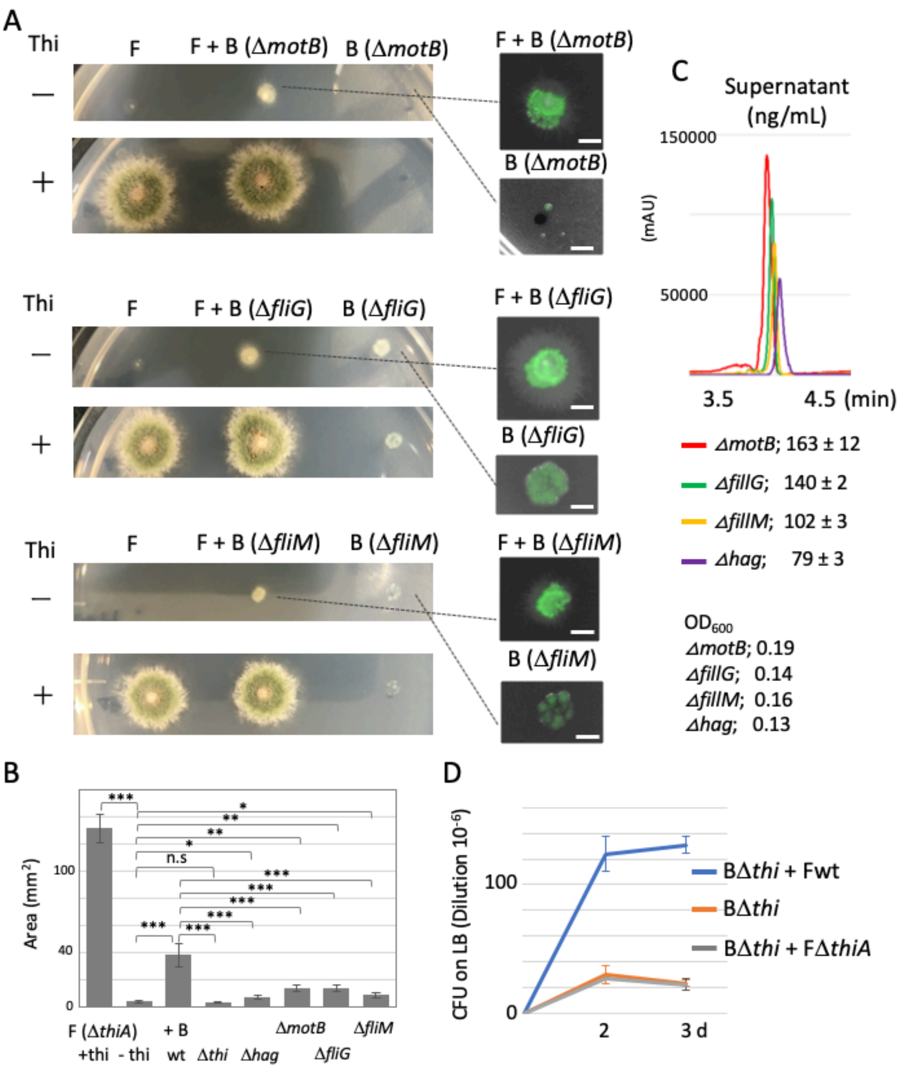

**Fig. S6. The growth defect of Δ*thiA* is recovered by co-culture with non-motile mutants.** (A) We co-culture an *A. nidulans* Δ*thiA* strain with three non-motile *B. subtilis* strains as follows. MotB; H+-coupled MotA-MotB flagellar stator. FliG; flagellar motor switch protein, physically transduces force from MotA to the rotation of FliF. FliM; flagellar motor switch protein, part of the basal body C-ring. The Δ*thiA* fungal colony shows a severe growth defect on the plate without thiamine, which is recovered by adding thiamine. The growth defect of Δ*thiA* is not recovered by co-culture with the non-motile mutants as co-culture with wild-type *B. subtilis*. Fungal colonies of *A. nidulans* (Δ*thiA*) mono-culture or co-cultivated with *B. subtilis* (Δ*motB*, Δ*fliG,* or Δ*fliM*) on minimal medium with/without thiamine grown for 2 days at 30°C. Each deletion strain expressing ZsGreen. Because flagella are required for bacterial dispersal on the hyphae, the non-motile mutants grow at the center of the fungal colony but do not reach the periphery of the fungal Δ*thiA* colony (a, right). (B) The area of fungal colonies is measured by ImageJ software. Error bar: S.D., n = 3, *** $P \leq 0.001$, * $P \leq 0.05$. (C) The amount of thiamine in supernatant in the mono-culture of *B. subtilis* non-motile mutants by LC-MS-MRM analysis. $OD_{600}$ are indicated. *B. subtilis* non-motile mutants synthesize and secrete thiamine in the medium. The growth effect on the fungal Δ*thiA* colony depend on the secretion amount in the four non-motile *B. subtilis* strains (B, C). (D) CFU on LB (dilution $10^{-6}$) in *B. subtilis* Δthi mono-culture, *B. subtilis* Δthi + *A. nidulans* wild-type co-culture, *B. subtilis* Δthi + *A. nidulans* Δ*thiA* co-culture in 200 ml minimal medium with 120 rpm shaking at 30°C for 2 and 3 days. CFU in *B. subtilis* Δthi + *A. nidulans* wild-type is higher than *B. subtilis* Δthi mono-culture, while that in *B. subtilis* Δthi + *A. nidulans* Δ*thiA* co-culture is comparable with *B. subtilis* Δthi mono-culture, suggesting a bi-directional thiamine transfer between *B. subtilis* and *A. nidulans*.

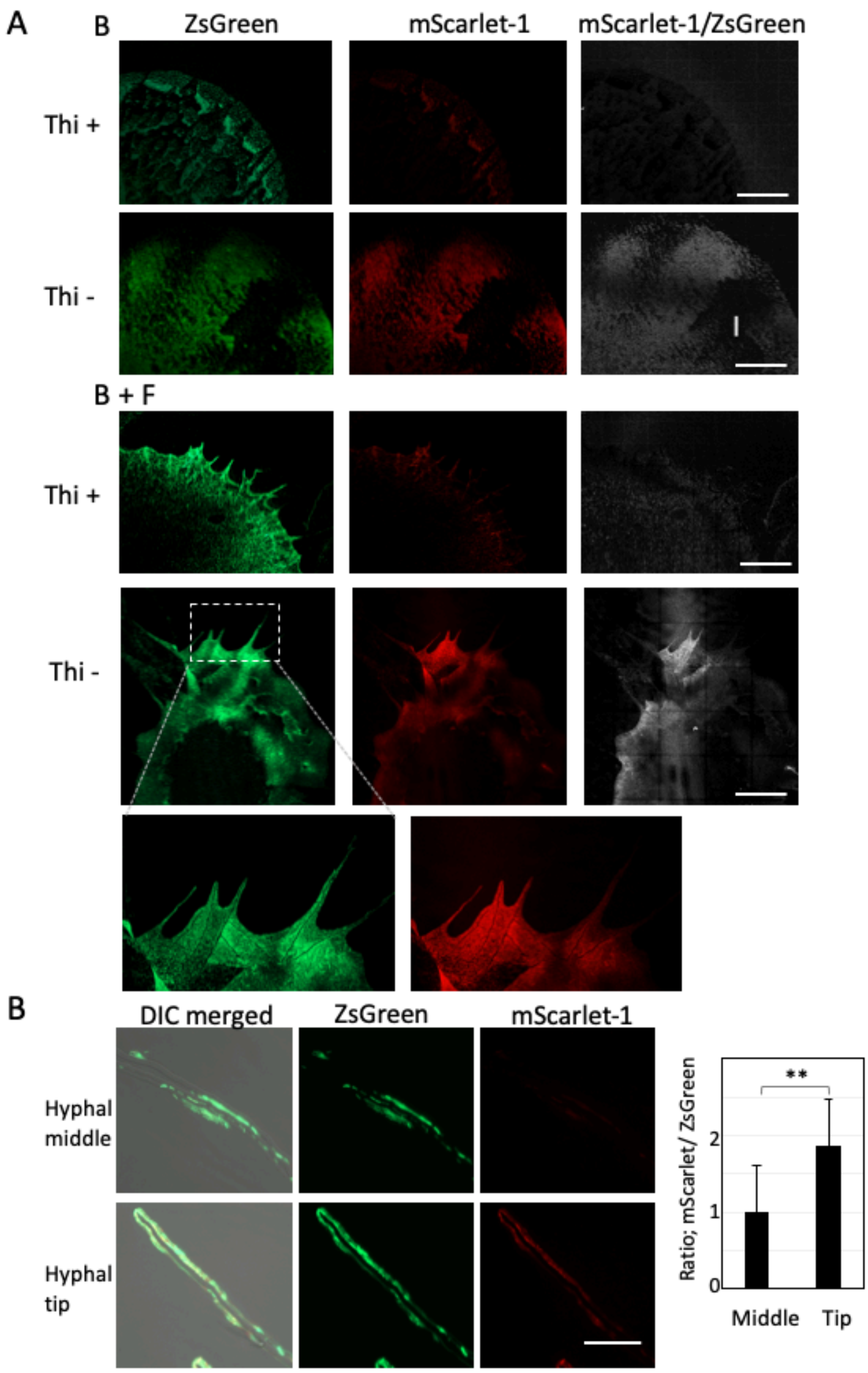

**Fig. S7.** ***B. subtilis*** **thiamine reporter strain**. (A) Colonies of the *B. subtilis* reporter strain in mono-culture and co-culture with *A. nidulans* on the minimal medium with/without thiamine grown for 2 days at 30°C. The images are constructed by 10 x 10 tiling of 500 x 500 µm image. Scale bar: 500 µm. (B) The *B. subtilis* reporter strain at hyphal middles and hyphal tips. Scale bar: 20 µm. Ratio of signal intensity, mScarlet-1/ZsGreen. Error bar: S.D., n = 75, ** P < 0.01.

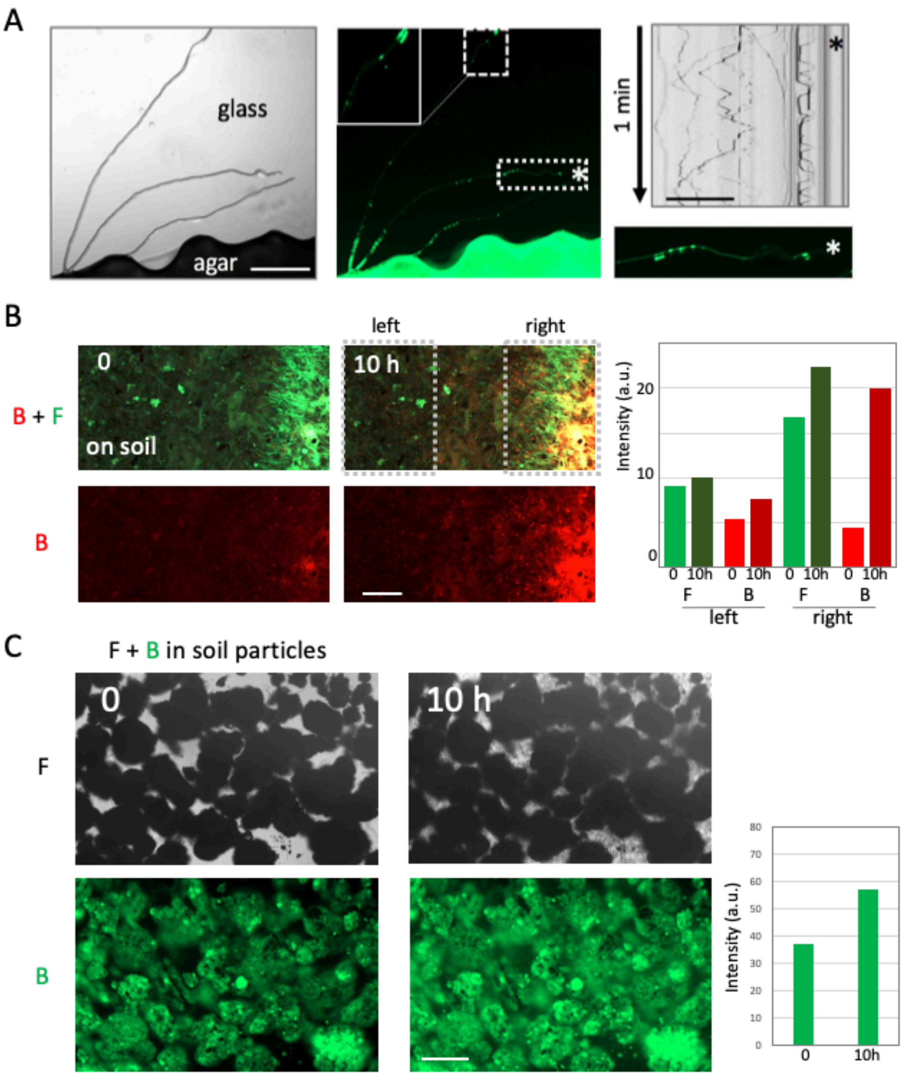

**Fig. S8. Bacterial movement along hyphae on glass surfaces and growth in soil.** (A) The point-inoculated *A. nidulans* and *B. subtilis* were grown on the agar media, cut out the 5 mm$^2$ piece and put on the cover glass sandwiching the fungus and bacteria between the glass and the agar block. Some hyphae continued to elongate on the glass surface to some extent without the agar media. Most bacteria stayed on the agar, however some bacteria cells still moved along the hyphae grown on the glass (Movie S11). *B. subtilis* (green) moved along the *A. nidulans* hyphae (DIC) grew on the glass surface from the agar block. Scale bar: 200 µm. Kymograph of *B. subtilis* movement along the hyphae from Movie S11. Total 1 min. Scale bar: 200 µm. (B) We cultivate *A. nidulans* and *B. subtilis* with soil and take time-lapse images for 10 h (Movie S12). The bacterial fluorescent signals were higher around the mycelium (right) as compared to the regions without mycelium (left), suggesting bacterial growth is sustained by the fungal network in soil, however it is difficult to track the fungal and bacterial growth in soil since it is opaque. Time-lapse images of *A. nidulans* hyphae (green) and *B. subtilis* (red) in soil at time 0 and 10 h from Video 12. Scale bar: 200 µm. Fluorescent signal intensity of *A. nidulans* (green) and *B. subtilis* (red) in left and right part at time 0 and 10 hours from Movie S12. (C) To monitor the fungal and bacterial growth and movement in soil, we used soil particles with sizes ~250 µm. Fungal growth is observed in gaps between soil particles, which is supplied with minimal media (1 µl mg$^{-1}$), over 10 hours. The signal intensity of bacteria labeled with green fluorescence also increased despite the high background fluorescence signal from soil particles. Time-lapse images of *A. nidulans* hyphae (DIC) and *B. subtilis* (green) in soil particles at time 0 and 10 h from Movie S13. Scale bar: 200 µm. Fluorescent signal intensity of *B. subtilis* (green) at time 0 and 10 hours.

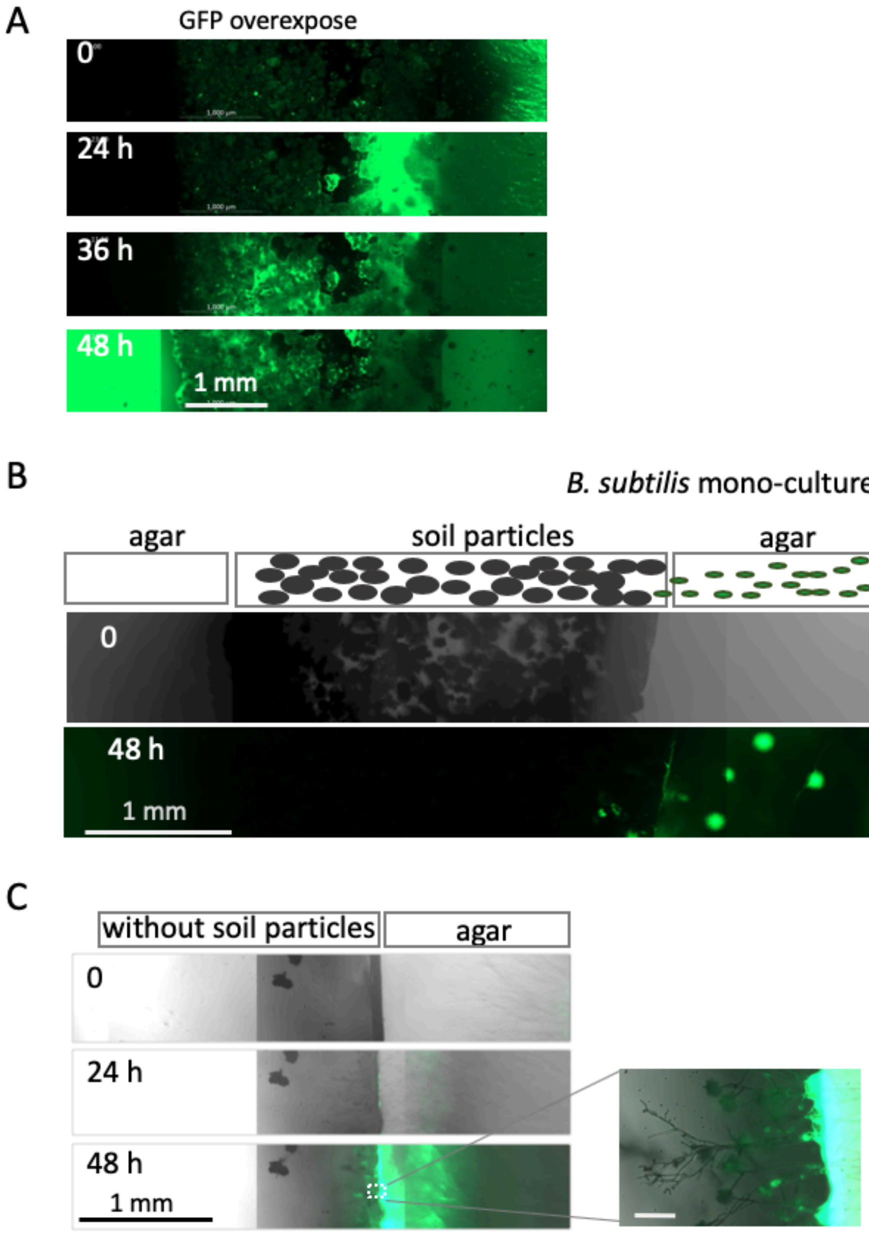

**Fig. S9. Bacterial migration on fungal colony in soil.** (A) Time-lapse images of bacterial migration (green) on growing hyphae in the soil particles sandwiched with two agar pieces at time 0, 24, 36 and 48 h from Movie S14. Scale bar: 1 mm. GFP overexposed in Fig. 5A. (B) Time-lapse images of bacterial migration (green) of *B. subtilis* mono-culture in soil particles sandwiched with two agar pieces at time 0 and 48 h from Movie S15. Scale bar: 1 mm. Without the fungus, however, after the bacteria reached the soil particles from the right agar slab, we observed no motion through the soil particles to the left agar slab. Instead they proliferated in the soil particles close to the right agar slab over 48 hours (C) Time-lapse images of bacterial migration (green) on growing hyphae without the soil particles sandwiched with two agar pieces at time 0, 24, and 48 h from Movie S16. Scale bar: 1 mm. Without soil particles, the fungal hyphae emerged from the right agar slab, extended to some extent and eventually stopped growing.

Table S1. *B. subtilis* genes differentially expressed in the co-culture condition[a]

| Gene name | **Product** | **Fold change in co-culture** |
|---|---|---|
| **Up regulated** | | |
| | **thiamine related** | |
| *thiU* | thiamine-binding protein | 4.69 |
| *thiC* | phosphomethylpyrimidine synthase ThiC | 4.68 |
| *thiV* | thiamine permease | 3.90 |
| *tenA* | thiaminase II | 3.68 |
| *thiX* | thiamine permease | 3.66 |
| *thiW* | cobalt ABC transporter | 3.57 |
| *thiF* | thiamine biosynthesis protein ThiS | 3.35 |
| *thiD* | bifunctional hydroxymethylpyrimidine kinase/phosphomethylpyrimidine kinase | 3.31 |
| *thiO* | glycine oxidase | 3.20 |
| *thiG* | thiazole synthase | 3.10 |
| *tenI* | thiamine phosphate synthase | 3.05 |
| *thiS* | thiamine biosynthesis protein ThiS | 2.65 |
| *thiT* | thiamine transporter ThiT | 2.01 |
| | **others** | |
| *glpD* | aerobic glycerol-3-phosphate dehydrogenase | 2.45 |
| *ydbN* | hypothetical protein | 2.36 |
| *cidA* | holin-like protein CidA | 2.21 |
| *ssuB* | aliphatic sulfonate ABC transporter ATP-binding protein | 2.12 |
| **Down regulated** | | |
| *pstC* | ABC transporter permease | -2.79 |
| *nrgA* | ammonium transporter NrgA | -2.64 |
| *ytnP* | MBL fold metallo-hydrolase | -2.11 |
| *pucR* | purine catabolism regulatory protein | -2.03 |

[a]Differentially expressed was defined by a fold change of >2 or <-2 and an RPKM of >10.

Table S2. *A. nidulans* down-regulated genes in co-culture with more than 8-fold

**Table S3. The down-regulated gene in co-culture with more than 8-fold**

| | | RPKM | | | |
|---|---|---|---|---|---|
| | FC (F_B / F) | Fungi | i_Bacteria | Gene name | info |
| AN2319 | -32.00 | 85.16 | 1.99 | | Ortholog of A. niger CBS 513.88 : An14g04210, A. oryzae RIB40 : AO090010000485, Aspergillus wentii : Aspwe1_0041701, Aspergillus sydowii : Aspsy1_0039595 and Aspergillus terreus NIH2624 : ATET_07606 |
| AN7159 | -28.18 | 43.31 | 0.78 | | Ortholog(s) have tripeptidyl-peptidase activity |
| AN4148 | -26.21 | 268.73 | 7.90 | xtrE | Putative xylose transporter; transcriptionally induced by growth on xylose |
| AN3427 | -24.00 | 29.16 | 0.25 | | Ortholog(s) have nucleolus, preribosome localization |
| AN5770 | -23.03 | 46.74 | 1.38 | | Ortholog of A. fumigatus Af293 : Afu6g06610, A. niger CBS 513.88 : An18g06470, A. oryzae RIB40 : AO090003000036, Aspergillus wentii : Aspwe1_0038379 and Aspergillus sydowii : Aspsy1_0149748 |
| AN6426 | -21.34 | 313.61 | 11.52 | | Has domain(s) with predicted metallopeptidase activity |
| AN0092 | -19.46 | 37.89 | 1.27 | | Putative methyltransferase with a predicted role in histidine metabolism |
| AN2578 | -15.99 | 109.69 | 5.23 | | Has domain(s) with predicted peroxidase activity |
| AN7832 | -15.41 | 44.80 | 2.24 | | Has domain(s) with predicted flavin adenine dinucleotide binding, oxidoreductase activity, acting on CH-OH group of donors activity and role in oxidation-reduction process |
| AN7711 | -14.81 | 49.37 | 2.62 | ndxB | Putative nudix hydrolase |
| AN3928 | -14.69 | 3758.72 | 294.90 | thiF | Putative thiazole synthase, enzyme of the thiamine biosynthesis pathway |
| AN8247 | -14.50 | 35.90 | 1.80 | | Ortholog(s) have role in maturation of 5.8S rRNA from tricistronic rRNA transcript (SSU-rRNA, 5.8S rRNA, LSU-rRNA), maturation of LSU-rRNA from tricistronic rRNA transcript (SSU-rRNA, 5.8S rRNA, LSU-rRNA) |
| AN9496 | -13.66 | 35.09 | 1.88 | | Has domain(s) with predicted cholestenol delta-isomerase activity, role in sterol metabolic process and endoplasmic reticulum, integral component of membrane localization |
| AN6621 | -13.59 | 104.00 | 5.84 | | Ortholog of A. nidulans FGSC A4 : AN7403, A. fumigatus Af293 : Afu8g01810, A. niger CBS 513.88 : An05g02280, A. oryzae RIB40 : AO090026000198 and Aspergillus wentii : Aspwe1_0151553, Aspwe1_0177691 |
| AN3206 | -13.35 | 195.46 | 11.70 | | Putative aryl-alcohol oxidase-related protein; protein expressed at decreased levels in a hapX mutant versus wild-type; transcript is induced by nitrate |
| AN0806 | -12.61 | 25.14 | 1.33 | | Ortholog(s) have role in single-species biofilm formation on inanimate substrate and cytosol, nucleus localization |
| AN5340 | -11.93 | 31.92 | 1.95 | | Ortholog(s) have role in endonucleolytic cleavage in 5'-ETS of tricistronic rRNA transcript (SSU-rRNA, 5.8S rRNA and LSU-rRNA), more |
| AN2292 | -11.73 | 27.76 | 1.71 | | Ortholog(s) have role in maturation of SSU-rRNA from tricistronic rRNA transcript (SSU-rRNA, 5.8S rRNA, LSU-rRNA) and cytoplasm, nucleolus localization |
| AN0956 | -11.69 | 35.78 | 2.33 | | Ortholog(s) have tRNA dihydrouridine synthase activity, role in tRNA modification and nucleus localization |
| AN9125 | -11.64 | 43.73 | 2.92 | | RNA polymerase III, subunit C34; locus contains the conserved upstream open reading frame (uORF) AN9125-uORF |
| AN3481 | -10.79 | 581.54 | 44.88 | | Ortholog(s) have carbohydrate binding activity |
| AN1949 | -10.74 | 69.84 | 5.02 | | Ortholog(s) have ATP-dependent RNA helicase activity, RNA binding activity |
| AN4178 | -10.46 | 104.64 | 7.69 | myg1 | UPF0160 domain-containing protein; intracellular; protein abundance decreased by menadione stress |
| AN5010 | -10.44 | 28.96 | 2.05 | | Ortholog(s) have role in cellular response to biotic stimulus, cellular response to starvation and filamentous growth of a population of unicellular organisms in response to biotic stimulus, more |
| AN2574 | -10.43 | 13.83 | 0.51 | | Has domain(s) with predicted UDP-N-acetylmuramate dehydrogenase activity, flavin adenine dinucleotide binding activity and role in oxidation-reduction process |
| AN5871 | -10.37 | 40.09 | 2.98 | | Ortholog(s) have rRNA (adenine) methyltransferase activity and role in rRNA methylation, ribosomal large subunit biogenesis |
| AN0946 | -10.22 | 51.83 | 3.95 | | Ortholog(s) have mRNA binding activity |
| AN4456 | -10.06 | 31.94 | 2.43 | | Ortholog(s) have role in cellular response to drug and small-subunit processome localization |
| AN4065 | -9.90 | 34.04 | 2.66 | | Ortholog of A. nidulans FGSC A4 : AN7503, A. fumigatus Af293 : Afu1g05410, Afu2g05420, A. niger CBS 513.88 : An11g09310, An18g04230 and A. oryzae RIB40 : AO090001000653, AO090009000402 |
| AN7671 | -9.58 | 37.33 | 3.00 | | Ortholog of A. fumigatus Af293 : Afu2g01440, A. niger CBS 513.88 : An03g05020, A. oryzae RIB40 : AO090701000815, Neosartorya fischeri NRRL 181 : NFIA_033740 and Aspergillus clavatus NRRL 1 : ACLA_093470 |
| AN9406 | -9.58 | 18.40 | 1.23 | trm1 | Putative 2,N2-dimethylguanosine tRNA methyltransferase; mutants are self-sterile |
| AN7630 | -9.57 | 69.20 | 5.50 | | Ortholog(s) have unfolded protein binding activity, role in ribosomal large subunit assembly and cytosol, cytosolic ribosome, preribosome, large subunit precursor localization |
| AN6865 | -9.31 | 42.61 | 3.51 | | Ortholog of A. fumigatus Af293 : Afu5g13120, A. niger CBS 513.88 : An14g06020, A. oryzae RIB40 : AO090120000437, Aspergillus wentii : Aspwe1_0170850 and Aspergillus sydowii : Aspsy1_0144687 |
| AN7142 | -9.19 | 207.66 | 18.69 | pacA | Putative secreted acid phosphatase with a predicted role in gluconic acid and gluconate metabolism |
| AN7177 | -9.18 | 134.80 | 11.69 | | Ortholog of A. fumigatus Af293 : Afu4g03322, A. niger CBS 513.88 : An14g01990, A. oryzae RIB40 : AO090011000139, Aspergillus sydowii : Aspsy1_0052594 and Aspergillus terreus NIH2624 : ATET_02136 |
| AN6934 | -9.12 | 15.11 | 0.97 | | Has domain(s) with predicted role in transmembrane transport and integral component of membrane localization |
| AN1524 | -8.83 | 158.82 | 14.34 | | Putative methylenetetrahydrofolate dehydrogenase (NAD+) with a predicted role in one-carbon metabolism |
| AN5884 | -8.69 | 121.68 | 10.96 | pyrF | Orotate phosphoribosyltransferase (OPRTase) that catalyzes the fifth step in pyrimidine biosynthesis |
| AN3573 | -8.68 | 20.65 | 1.71 | | Has domain(s) with predicted oxidoreductase activity, acting on the aldehyde or oxo group of donors, NAD or NADP as acceptor activity and role in oxidation-reduction process |
| AN0072 | -8.66 | 34.73 | 3.08 | | Putative pseudouridine synthase with a predicted role in pyrimidine metabolism |
| AN0259 | -8.66 | 40.18 | 3.56 | | Putative adenylate kinase with a predicted role in nucleotide salvage pathways |
| AN0272 | -8.63 | 16.16 | 1.20 | | Ortholog of A. fumigatus Af293 : Afu1g02880, A. niger CBS 513.88 : An01g05030, A. oryzae RIB40 : AO090005000779, Aspergillus wentii : Aspwe1_0025563 and Aspergillus sydowii : Aspsy1_0055099 |
| AN0796 | -8.63 | 27.96 | 2.52 | | Has domain(s) with predicted role in rRNA processing and small-subunit processome localization |
| AN0204 | -8.63 | 16.97 | 1.31 | | Has domain(s) with predicted ATP binding, ATP-dependent helicase activity, nucleic acid binding activity |
| AN3706 | -8.59 | 2894.33 | 315.05 | | S. cerevisiae ortholog RPS10A has role in rRNA export from nucleus; palA-dependent expression independent of pH |
| AN5713 | -8.58 | 61.38 | 5.45 | cct7 | Putative chaperonin complex component, TCP-1 eta subunit; ortholog of S. cerevisiae Cct7p; expression reduced after exposure to farnesol |
| AN4102 | -8.57 | 49.23 | 4.44 | bglA | Putative beta-glucosidase; induced by carbon starvation-induced autophagy |
| AN4346 | -8.49 | 102.48 | 9.29 | | Has domain(s) with predicted nucleic acid binding, nucleotide binding activity |
| AN5702 | -8.41 | 52.95 | 4.83 | | Ortholog(s) have role in ribosomal large subunit biogenesis, ribosomal small subunit biogenesis and cytosolic large ribosomal subunit, nucleolus localization |
| AN8799 | -8.23 | 13.98 | 1.01 | | Has domain(s) with predicted tRNA (adenine-N1-)-methyltransferase activity, role in tRNA methylation and tRNA (m1A) methyltransferase complex localization |
| AN7214 | -8.23 | 37.65 | 3.49 | | Ortholog of A. nidulans FGSC A4 : AN8162, A. fumigatus Af293 : Afu5g01210, A. niger CBS 513.88 : An16g00660 and A. oryzae RIB40 : AO090102000383, AO090120000014, AO090001000406, AO090103000238 |
| AN0173 | -8.09 | 14.24 | 1.08 | | Ortholog of A. niger CBS 513.88 : An01g02440, A. oryzae RIB40 : AO090026000850, Aspergillus wentii : Aspwe1_0442639, Aspergillus sydowii : Aspsy1_0027713 and Aspergillus terreus NIH2624 : ATET_07718 |
| AN3176 | -8.04 | 18.36 | 1.60 | spb4 | ATP-dependent rRNA helicase |
| AN8009.2 | -7.47 | 9914.37 | 1315.89 | nmt1 | Ortholog of A. fumigatus nmt1 and S. cerevisiae THI13; repressed by starvation-induced autophagy; thi ORF\|Unchar ChrII_A_nidulans_FGSC_A4 |

Table S3. *A. nidulans* up-regulated genes in co-culture with more than 8-fold

**Table S4. The up-regulated gene in co-culture with more than 8-fold**

| | FC (F_B / F) | RPKM Fungi | RPKM Fungi_Bacteria | Gene name | info |
|---|---|---|---|---|---|
| AN5488 | **33.86** | 7.23 | 248.27 | | Protein of unknown function |
| AN6862 | **25.57** | 33.64 | 678.67 | | Ortholog of A. niger CBS 513.88 : An03g01640, Neosartorya fischeri NRRL 181 : NFIA_074040, Aspergillus wentii : Aspwe1_0170839 and Aspergillus versicolor : Aspve1_0035273 |
| AN7735 | **17.66** | 15.11 | 243.64 | eglD | Putative endoglucanase; protein localized to the conidial cell wall |
| AN7217 | **16.40** | 58.66 | 761.23 | | Protein of unknown function |
| AN9108 | **15.78** | 13.72 | 198.11 | | Has domain(s) with predicted heme binding activity |
| AN7378 | **15.50** | 44.37 | 588.52 | | Ortholog of A. niger CBS 513.88 : An15g07560, A. oryzae RIB40 : AO090103000092, Neosartorya fischeri NRRL 181 : NFIA_102090 and Aspergillus versicolor : Aspve1_0187655 |
| AN5083 | **14.66** | 10.93 | 148.92 | | Has domain(s) with predicted hydrogen ion transmembrane transporter activity, role in ATP hydrolysis coupled proton transport and vacuolar proton-transporting V-type ATPase, V0 domain localization |
| AN5015 | **14.48** | 132.88 | 1495.55 | conJ | Putative conidiation gene; transcript induced by light in in developmentally competent mycelia; double conF and conJ deletion results in increased cellular glycerol or erythritol leading to delayed germination and desiccation resistance |
| AN8043 | **14.17** | 134.44 | 1481.25 | | Protein of unknown function |
| AN8333 | **14.06** | 79.87 | 875.22 | phiA | Protein required for normal phialide development; expression increased in the presence of bafilomycin B1 and concanamycin A; PhiA localizes to conidia and phialides |
| AN7799 | **13.78** | 0.88 | 27.52 | | Protein expressed at increased levels during osmoadaptation; short-chain alcohol dehydrogenase domain |
| AN5490 | **13.42** | 6.45 | 88.11 | | Ortholog of A. niger CBS 513.88 : An08g10190, A. oryzae RIB40 : AO090003000446, Aspergillus versicolor : Aspve1_0053274, Aspergillus niger ATCC 1015 : 37923-mRNA and Aspergillus sydowii : Aspsy1_0150005 |
| AN2617 | **13.33** | 0.50 | 21.62 | | Protein of unknown function |
| AN7102 | **12.66** | 116.86 | 1140.28 | | Ortholog of A. fumigatus Af293 : Afu4g03830, A. niger CBS 513.88 : An14g01068, A. oryzae RIB40 : AO090011000343, Aspergillus wentii : Aspwe1_0030332 and Aspergillus sydowii : Aspsy1_0060599 |
| AN0483 | **12.11** | 68.29 | 665.34 | | Ortholog of Neosartorya fischeri NRRL 181 : NFIA_058180, Aspergillus versicolor : Aspve1_0158177, Aspergillus clavatus NRRL 1 : ACLA_085420 and Aspergillus zonatus : Aspzo1_0018005 |
| AN4935 | **11.72** | 212.68 | 2159.53 | ran1 | Ortholog(s) have protein serine/threonine kinase activity |
| AN8154 | **11.15** | 0.21 | 13.81 | | Protein of unknown function |
| AN7945 | **10.97** | 3.53 | 46.73 | hk2 | Putative histidine-containing phosphotransfer protein |
| AN9365 | **10.72** | 6.19 | 67.98 | xlnB | Protein with endo-1,4-beta-xylanase activity, involved in degradation of xylans |
| AN3308 | **10.53** | 35.34 | 328.37 | amyD | Putative alpha-amylase with a predicted role in starch metabolism; predicted glycosyl phosphatidylinositol (GPI)-anchor |
| AN3680 | **10.49** | 442.33 | 5274.88 | | Ortholog of A. fumigatus Af293 : Afu4g12500, A. niger CBS 513.88 : An01g07980, Aspergillus wentii : Aspwe1_0033896 and Aspergillus zonatus : Aspzo1_0129782 |
| AN8930 | **10.46** | 5.94 | 64.45 | | Has domain(s) with predicted DNA binding, RNA polymerase II transcription factor activity, sequence-specific DNA binding, zinc ion binding activity and role in regulation of transcription, DNA-templated, transcription, DNA-templated |
| AN0397 | **10.27** | 3.63 | 44.58 | | Protein of unknown function |
| AN8132 | **10.16** | 1.72 | 27.89 | | Has domain(s) with predicted metallopeptidase activity, zinc ion binding activity and role in proteolysis |
| AN4722 | **10.11** | 167.32 | 1326.93 | | Predicted glycosylphosphatidylinositol (GPI)-anchored protein |
| AN1740 | **9.99** | 0.96 | 20.35 | | Protein of unknown function |
| AN6653 | **9.98** | 37.30 | 330.14 | acuE | Malate synthase, required for utilization of acetate as carbon source; transcription induction by acetate mediated by FacB; carbon catabolite repression mediated by CreA; transcription induction by long-chain fatty acids mediated by FarA |
| AN1826 | **9.85** | 23.96 | 213.16 | | Has domain(s) with predicted hydrolase activity |
| AN5553 | **9.74** | 7.70 | 73.46 | CYP5080D1 | Putative cytochrome P450 |
| AN6642 | **9.72** | 21.56 | 187.14 | enaA | Putative potassium-transporting ATPase with a predicted role in energy metabolism; locus contains the conserved upstream open reading frames (uORFs) AN6642-uORF and AN6642-uORF.2 |
| AN0950 | **9.58** | 230.22 | 1810.96 | | Ortholog of Aspergillus versicolor : Aspve1_0036555 and Aspergillus sydowii : Aspsy1_0038433 |
| AN8966 | **9.51** | 50.02 | 421.64 | | Has domain(s) with predicted neurotransmitter:sodium symporter activity, role in neurotransmitter transport and integral component of membrane localization |
| AN7215 | **9.48** | 0.72 | 16.85 | | Protein of unknown function |
| AN6274 | **9.32** | 30.26 | 251.27 | | Has domain(s) with predicted oxidoreductase activity and role in metabolic process |
| AN1731 | **9.29** | 8.13 | 73.25 | prnD | Putative proline dehydrogenase with a predicted role in proline metabolism; expression is regulated by carbon and nitrogen repression; negatively regulated by CreA |
| AN4277 | **9.23** | 7.67 | 69.27 | | Ortholog(s) have glucose transmembrane transporter activity, role in cellular response to drug, glucose transport and plasma membrane localization |
| AN6881 | **9.23** | 18.52 | 151.96 | | Ortholog of A. fumigatus Af293 : Afu8g07160, A. niger CBS 513.88 : An03g01630, Neosartorya fischeri NRRL 181 : NFIA_099950 and Aspergillus versicolor : Aspve1_0035247 |
| AN3310 | **8.80** | 138.85 | 946.32 | | Predicted glycosylphosphatidylinositol (GPI)-anchored protein |
| AN8436 | **8.79** | 1.06 | 18.43 | | Has domain(s) with predicted hydrolase activity, acting on glycosyl bonds activity |
| AN5086 | **8.72** | 5.32 | 49.38 | SpoC1-C1D | Conidium-specific RNA of unknown function; member of the SpoC1 gene cluster |
| AN5942 | **8.67** | 17.26 | 133.84 | | Ortholog of Aspergillus versicolor : Aspve1_0148858 and Aspergillus sydowii : Aspsy1_0029071 |
| AN2817 | **8.62** | 9.07 | 74.37 | | Protein of unknown function |
| AN8077 | **8.54** | 2.42 | 27.98 | | Has domain(s) with predicted role in transmembrane transport and integral component of membrane localization |
| AN7990 | **8.48** | 48.92 | 372.27 | | Ortholog of A. niger CBS 513.88 : An11g06780, Aspergillus wentii : Aspwe1_0176973, Aspergillus sydowii : Aspsy1_0047960 and Aspergillus terreus NIH2624 : ATET_06661 |
| AN9377 | **8.31** | 9.87 | 76.15 | | Ortholog of A. fumigatus Af293 : Afu3g02940, A. niger CBS 513.88 : An01g00550, A. oryzae RIB40 : AO090120000006, Aspergillus wentii : Aspwe1_0112345 and Aspergillus sydowii : Aspsy1_0163833 |
| AN8627 | **8.30** | 16.01 | 119.82 | | Has domain(s) with predicted transferase activity, transferring glycosyl groups activity and membrane localization |
| AN3522 | **8.19** | 0.66 | 13.89 | fhbB | Flavohemoprotein, involved (with FhbA) in sterigmatocystin production |
| AN5489 | **8.18** | 10.75 | 80.76 | | Ortholog of A. fumigatus Af293 : Afu6g13220, A. niger CBS 513.88 : An08g10180, A. oryzae RIB40 : AO090003000447, Aspergillus wentii : Aspwe1_0166336 and Aspergillus sydowii : Aspsy1_0886379 |
| AN0773 | **8.09** | 35.99 | 257.50 | | Has domain(s) with predicted oxidoreductase activity and role in oxidation-reduction process |
| AN6084 | **8.07** | 35.77 | 254.49 | | Ortholog of A. fumigatus Af293 : Afu2g09220, A. niger CBS 513.88 : An12g04880, A. oryzae RIB40 : AO090011000773, Aspergillus wentii : Aspwe1_0678682 and Aspergillus sydowii : Aspsy1_0152386 |

Table S4. strains used in this study

| Strain | Genotype | Source |
| --- | --- | --- |
| *A. nidulans* | | |
| TN02A3 | *pyrG89; argB2; ΔnkuA::argB; pyroA4* | (*1*) |
| Δ*thiA* | *biA1*; *argB2*; Δ*thiA::argB* | (*2*) |
| *B. subtilis 3610* | Wild type | (3) |
| *B. subtilis 168* | *trpC2* | Lab stock |
| Δ*hag* | *168* Δ*hag::Cm*$^{R}$ | This study |
| Δ*thi-operon* | *168* Δ*thi-operon::Spec*$^{R}$ | This study |
| thi-rep | *3610* + pHY300MK<br>(Pveg-ZsGreen, PtenA-mScarlet-1:: *Tet*$^{R}$) | This study |

Table S5. Nucleotides used in this study

| Primer name | Sequence (5' to 3') |
|---|---|
| tenA-5 | GTAGGATCACCGGAGCATGAA |
| tenA-N5 | GCTTCGCGCACTGATTGAA |
| tenA-N3 | ctttatccaattttcGAAAATAAAAAAACCACTTTCCCGCAA |
| tenA-C5 | tcactaacctgccccGATCACCAGCTGATGGTGA |
| tenA-C3 | GGACGTGCCGCTTTGACAACA |
| tenA-3 | CACAAGCTCTCCGCCAAGGTA |
| Cm-Fw | GGTTTTTTTATTTTCgaaaattggataaagtgggatatttttaaaatatatattta |
| Cm-Rv | ATCACCATCAGCTGGTGATCggggcaggttagtgacatta |
| hag N5 | AGAGCCATTTGAAAAGTCTACTGC |
| hag-N3 | AGATCTCCATATAATTTTTGTGTTTTGTTCCTCCCTGAAT |
| spc-Fw | ATTCAGGGAGGAACAAAACACAAAAATTATATGGAGATCT |
| spc-Rv | CGCCAAGGTCTTTTTTAAAAAAGCTTCACTAAATTAAAGT |
| hag-C5 | ACTTTAATTTAGTGAAGCTTTTTTAAAAAAGACCTTGGCG |
| hag-C3 | AGACCTGTTATTCTTGTGACCATC |
| hag-3 | CTACAAATAACCCAAGAAATTCAG |
| tenA-5 | GTAGGATCACCGGAGCATGAA |
| tenA-mSca-Rv | TTCACCTTTAGATACcacaaatcattccccctctg |

**Video Legends**

Movie S1. *B. subtilis* (green) movement along *A. nidulans* hyphae (DIC in the first image) on the minimum agar plate. The sequence of three movies, 5 images/s, total 30 s, scale bar: 50 μm.

Movie S2. *B. subtilis* (green) movement along *A. nidulans* hyphae (DIC in the first image) on the minimum agar media. 20 images/s, total 30 s, scale bar: 20 μm.

Movie S3. *B. subtilis* (red) movement along *A. nidulans* hyphae (green) on the minimum agar plate. 12 images/s, total 30 s, scale bar: 25 μm.

Movie S4. *A. nidulans* hyphae (DIC, left) surrounded by moving *B. subtilis* (green, right) on the minimum agar plate. 5 images/s, total 20 s, scale bar: 50 μm.

Movie S5. *B. subtilis* flagella mutant (Δ*hag*) flow along *A. nidulans* hyphae (DIC in the first image). Every min, total 1 h, scale bar: 20 μm.

Movie S6. Dispersion of *B. subtilis* (168, green) on growing *A. nidulans* colony (DIC). Every 10 min, total 15 h, Scale bar: 100 μm.

Movie S7. Colony extension of *B. subtilis* (168, green) mono-culture. Every 10 min, total 18 h, Scale bar: 100 μm.

Movie S8. Proliferation of *B. subtilis* flagella mutant (Δ*hag,* green) and *A. nidulans* (DIC). Every 10 min, total 14 h, Scale bar: 100 μm.

Movie S9. Dispersion of *B. subtilis* (red) on growing *A. nidulans* mycelium (green). Every 10 min, total 19 h, Scale bar: 200 μm.

Movie S10. Dispersion of *B. subtilis* (green) on growing *A. nidulans* mycelium network (DIC) in 17 h from the *B. subtilis* inoculation. Every 10 min, total 20 h, Scale bar: 100 μm.

Movie S11. *B. subtilis* (green) movement along *A. nidulans* hyphae (DIC in the first image) on the glass surface. 10 images/s, total 1 min, scale bar: 100 μm.

Movie S12. Proliferation of *B. subtilis* (red) and *A. nidulans* (green) in the soil. Every 10 min, total 10 h, Scale bar: 200 μm.

Movie S13. Proliferation of *B. subtilis* (green) and *A. nidulans* (DIC) in the soil particles. Every 30 min, total 13 h, Scale bar: 200 μm.

Movie S14. Migration of *B. subtilis* colony (green) on mycelium extension of *A. nidulans* (DIC) from the right agar, through the soil particles, to the left agar. Every 30 min, total 48 h, Scale bar: 1000 μm.

Movie S15. No migration of *B. subtilis* colony (green) without *A. nidulans* from the right agar to soil particles. Every 30 min, total 48 h, Scale bar: 1000 μm.

Movie S16. Proliferation of *B. subtilis* colony (green) and *A. nidulans* colony (DIC) from the right agar without soil particles. Every 30 min, total 48 h, Scale bar: 1000 μm.

Movie S17. *Pantoea* sp. cells movement along *Trichoderma* sp. hyphae on the minimum agar plate. 3 images/s, total 30 s, scale bar: 10 μm.

**Refences**

1. Nayak T, et al. (2006) A versatile and efficient gene-targeting system for *Aspergillus nidulans*. Genetics 172: 1557-66.
2. 23. Shimizu M, et al. (2016) Thiamine synthesis regulates the fermentation mechanisms in the fungus *Aspergillus nidulans*. Biosci. Biotechnol. Biochem. 80: 1768-1775.
3. Branda SS, et al. (2001) Fruiting body formation by *Bacillus subtilis*. Proc. Natl. Acad. Sci. U.S.A. 98: 11621-6.
